# *Ab initio* prediction of RNA structure ensembles with RNAnneal

**DOI:** 10.64898/2026.02.01.703098

**Authors:** Lukas Herron, Yunrui Qiu, Anjali Verma, Venkata Sai Sreyas Adury, Richard John, Suemin Lee, Shams Mehdi, Disha Sanwal, John S. Schneekloth, Pratyush Tiwary

## Abstract

RNA utilizes three-dimensional structure in addition to sequence to carry out diverse functions in gene expression and disease. Much like well-folded proteins, RNAs adopt specific three-dimensional structures to carry out their function. Yet comparatively few RNA structures have been solved by atomic resolution structural techniques, in part because unlike structured proteins, RNAs fold into heterogeneous ensembles of interconverting structures that pose a challenge for high-resolution structure probing methods. In this work, we introduce RNAnneal as a method for RNA structural ensemble prediction that seamlessly integrates generative deep learning with statistical physics and molecular dynamics modeling. Given the primary sequence, RNAnneal uses *ab inito* (i.e., first principles) modeling to sample an ensemble of 3D structures, which, in turn, are used to train an ensemble of unsupervised deep learning models. The RNAnneal score, representing the consensus of the deep learning models, is then used to evaluate the 3D structures. We evaluated RNAnneal structures against 16 experimentally-resolved conformations (ERCs) of riboswitch RNAs and found that pseudoknot-free (PK-free) ERCs were well-reproduced by RNAnneal, with clear avenues for improving performance even on PK-comprising structures. Furthermore, we found that the RNAnneal score outperforms the Rosetta score and a state-of-the-art RNA forcefield on the task of classifying ERCs from decoys. We then introduce the interaction entropy as a measure of conformational heterogeneity within an ensemble and use it to assess our predictions. RNAnneal thus provides a generalizable framework for predicting RNA structural ensembles that will accelerate RNA-targeted drug discovery and the design of functional RNA molecules.

---

RNA was discovered to be the messenger of genetic information, carrying the code for polypeptide sequences from the nucleus to the ribosome, in 1961.^1^ Six decades later, our understanding of RNA has expanded to include noncoding, molecular recognition, and enzymatic functions, establishing the transcriptome as a complex web of functional, interacting RNA molecules that orchestrate gene expression and disease at multiple levels.^2^ Interrogation of the transcriptome continues to yield insights that are critical for understanding basic biology and designing next-generation therapeutics.^3,4^ The approval of small molecules such as risdiplam^5^ and successes of mRNA vaccines^6,7^ highlight the role of RNA as both a target and therapeutic modality.

Understanding how RNA sequence gives rise to three-dimensional structure and function remains a frontier for theoretical and computational science. Accurate structure models would inform decision making in RNA therapeutic development in a similar manner to proteins, where atomic resolution structure models are frequently considered when evaluating the therapeutic potential of protein targets. These models, however, are rarely available for RNAs. To illustrate, approximately 100 RNA-only structures are deposited in the Protein Data Bank (PDB) annually compared to 10,000 for proteins.^8^

Furthermore, most PDB entries are solved by high-resolution methods like nuclear magnetic resonance spectroscopy (NMR), X-ray crystallography, or cryo-electron microscopy. These methods operate by detecting an average signal from the conformational ensemble to generate a structural model, and are therefore most accurate when the ensemble is homogeneous. In the case of less ordered structures, the detector receives a superposition of the individual conformations’ signals that can be difficult to deconvolve. The structural heterogeneity of RNAs therefore presents a severe limitation to such ensemble averaged methods. New experimental and computational methods are required to accurately characterize such heterogeneous conformational ensembles.^9–11^ Here, we introduce RNAnneal as a computational method for predicting RNA structural ensembles. RNAnneal integrates modeling principles from generative deep learning, statistical physics, and molecular simulations into an *ab inito* conformational sampler and a deep learning score model.

## RNAnneal

RNAnneal is an unsupervised, *ab initio* method for predicting RNA structural ensembles that integrates generative deep learning with statistical physics and Molecular Dynamics (MD). RNAnneal approaches structure prediction in the spirit of a needle-in-a-haystack classification problem: a candidate structural ensemble is generated by broadly sampling secondary structures, assembling them into 3D models, selecting representatives, and performing additional conformational sampling with MD simulations (Fig. 1a). If the candidate ensemble contains experimentally resolved conformations (ERCs), then structure prediction amounts to picking them out of the ensemble. To this end, an RNAnneal score is developed based on a physics-inspired generative model, and used to rank candidate structures (Fig. 1b-e). The RNAnneal score is parameterized from the candidate conformations by an ensemble of Thermodynamic Maps, a type of generative model that infers the dependence of the conformational ensemble on environmental variables such as temperature or pressure (Fig. 1b-c). Thermodynamic maps are probabilistic generative models based on the denoising diffusion framework^13^, and have been shown to accurately predict molecular equilibrium distributions from sparse data collected across different thermodynamic conditions.^12,14,15^ Consistent with these works, one may tailor the RNAnneal score to a user-specified temperature by annealing the ensemble of score models.

**FIG. 1.**
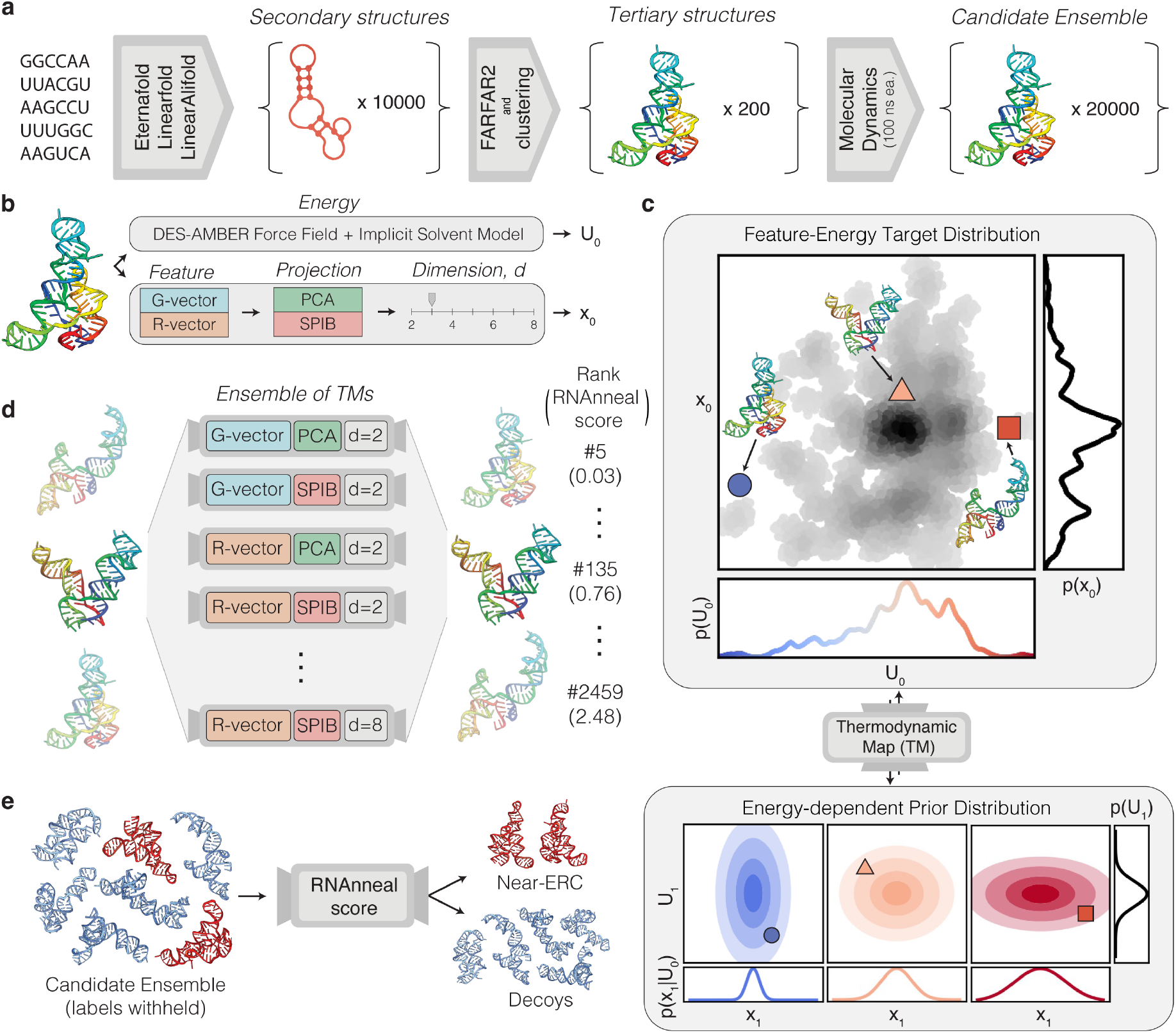
Illustrative overview of RNAnneal. **a**. The conformational sampling component of RNAnneal is illustrated. Starting with the primary sequence, a large number of secondary and 3D structures are predicted in succession. The 3D conformations are clustered into representatives before each is simulated for 100ns using implicit solvent molecular dynamics. Together, the simulation frames form a set of diverse candidate conformations that will be scored by RNAnneal. **b**. Energetic features (upper track) and latent representations (lower track) are extracted from each conformation. The latent features are constructed by projecting geometric descriptors (G- and R-vectors) into a reduced space of dimension *d* using Principal Component Analysis (PCA) and a State Predictive Information Bottleneck (SPIB) model. **c**. The structure of a thermodynamic map^12^ (TM) is illustrated: latent representations, **x**_0_ are mapped onto an energy-dependent prior distribution, while the energies, *U*_0_ themselves are mapped onto a unit normal distribution. **d**. TM models are trained on geometric features of dimension *d* produced by PCA and SPIB. The RNAnneal score is derived from the log-likelihood of the candidate conformations under the models. **e**. The RNAnneal score is suited for the task of classifying experimentally resolved conformations (ERCs) from decoys.

An essential quality of any RNA structure prediction method is the ability to generalize beyond ERCs, which form the set of known RNA structures. In computational modeling, ERCs are experimentally validated structure models that are serve as ground-truths for computational structure prediction methods. Most approaches are explicitly parameterized, trained, and validated against these experimentally determined structures. However, ERCs capture only a subset of the full conformational ensemble, and assessing model generalization beyond ERCs is critical yet challenging because the true structural ensembles are unknown. Unlike supervised methods that require labeled training data, RNAnneal learns a score function de novo for each sequence, avoiding issues of overfitting to previously observed structures. Within this framework, one may evaluate the score model on ERCs and examine if the model selects for them consistently. If they are well predicted, then the highest-scoring decoy structures may be reconsidered as potential alternative conformations.

We compared conformations sampled by RNAnneal to validated ERCs for 16 riboswitches and found agreement in their interaction compositions, measured by relative fractions of stacking, Watson-Crick and non-Watson-Crick contacts. (Fig. 2a-b). After simulating the ERCs for 100ns, we found their structures to be dynamic. This informed our decision to use the interaction network fidelity (INF) and root mean squared deviation (RMSD) together as complementary evaluation metrics. RMSD captures global geometric deviation but is sensitive to alignment, while INF captures interaction fidelity independent of alignment (Fig. 2c-f). In light of the observed flexibility, we evaluated RNAnneal against the simulated ERCs and found that pseudoknot-free (PK-free) near-ERCs were consistently sampled. Following this evaluation, we demonstrate that the unsupervised RNAnneal score prioritizes ERCs over decoys. For each RNA, 20,000 structures (including the ERC) were sampled, and RNAnneal was employed to select ten representative structures for each of the riboswitches. For 11/16 riboswitches, a representative with RMSD *<* 10Å or INF *>* 0.75 from the ERC was selected. Lastly, we introduce the interaction entropy as a per-nucleotide measure of structural heterogeneity within an ensemble (Fig. 3a-b).

**FIG. 2.**
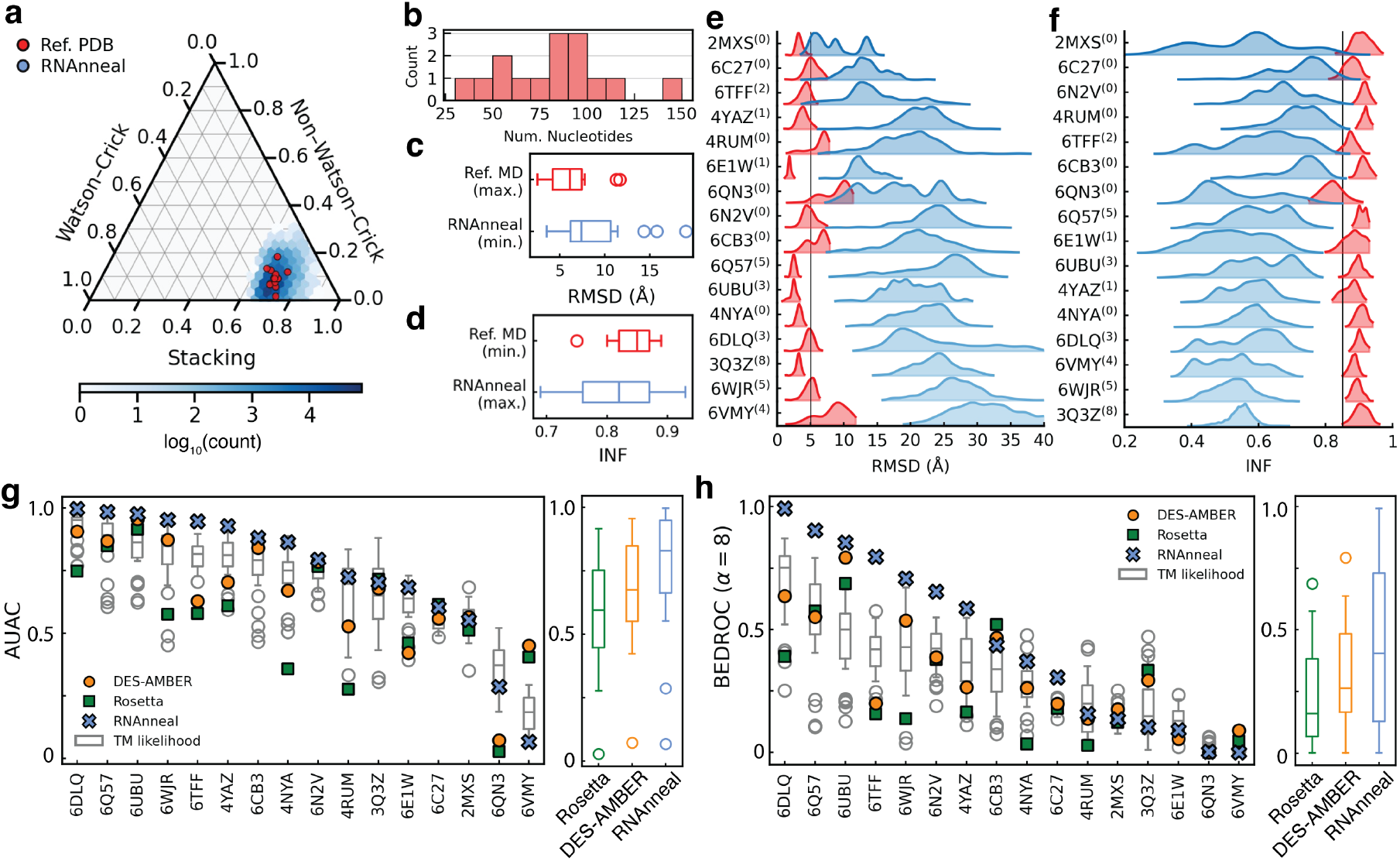
RNAnneal evaluated on the riboswitch dataset. **a**. Comparison between the interaction composition of ERC (red) and RNAnneal conformations (blue). **b**. Distribution of sequence lengths for the riboswitch set. **c**. Maximum RMSD values from MD simulations of the ERC are compared to the minimum RMSD among the RNAnneal conformations. (Median [IQR]; Ref. MD: 6.1Å [4.2 − 7.4Å]; RNAnneal: 7.4Å [6.2 − 10.7Å]). **d**. Minimum INF values from MD simulations of the ERC are compared to the maximum INF among the RNAnneal conformations (Median [IQR]; Ref. MD: 0.85 [0.83 − 0.87]; RNAnneal: 0.80 [0.76 − 0.87]). **e**. RMSD comparison between simulations of the ERC and conformations predicted by RNAnneal. **f**. INF comparison between simulations of the ERC and conformations predicted by RNAnneal. Conformations with INF ≥ 0.85 (vertical line) are classified as ERCs when evaluating the RNAnneal score (g and h). The superscripts denote the number of PK base pairs in the reference conformations (e and f).**g**. Comparison of AUAC values on the ERC classification task (Median (IQR); Rosetta: 0.59 [0.45 − 0.75]; DES-AMBER: 0.67 [0.55 − 0.85]; RNAnneal: 0.83 [0.66 − 0.95]). **h**. Comparison of BEDROC values for the ERC classification task (Median (IQR); Rosetta: 0.16 [0.07 − 0.38]; DES-AMBER: 0.26 [0.17 − 0.48]. RNAnneal: 0.40 [0.13 − 0.73]).

**FIG. 3.**
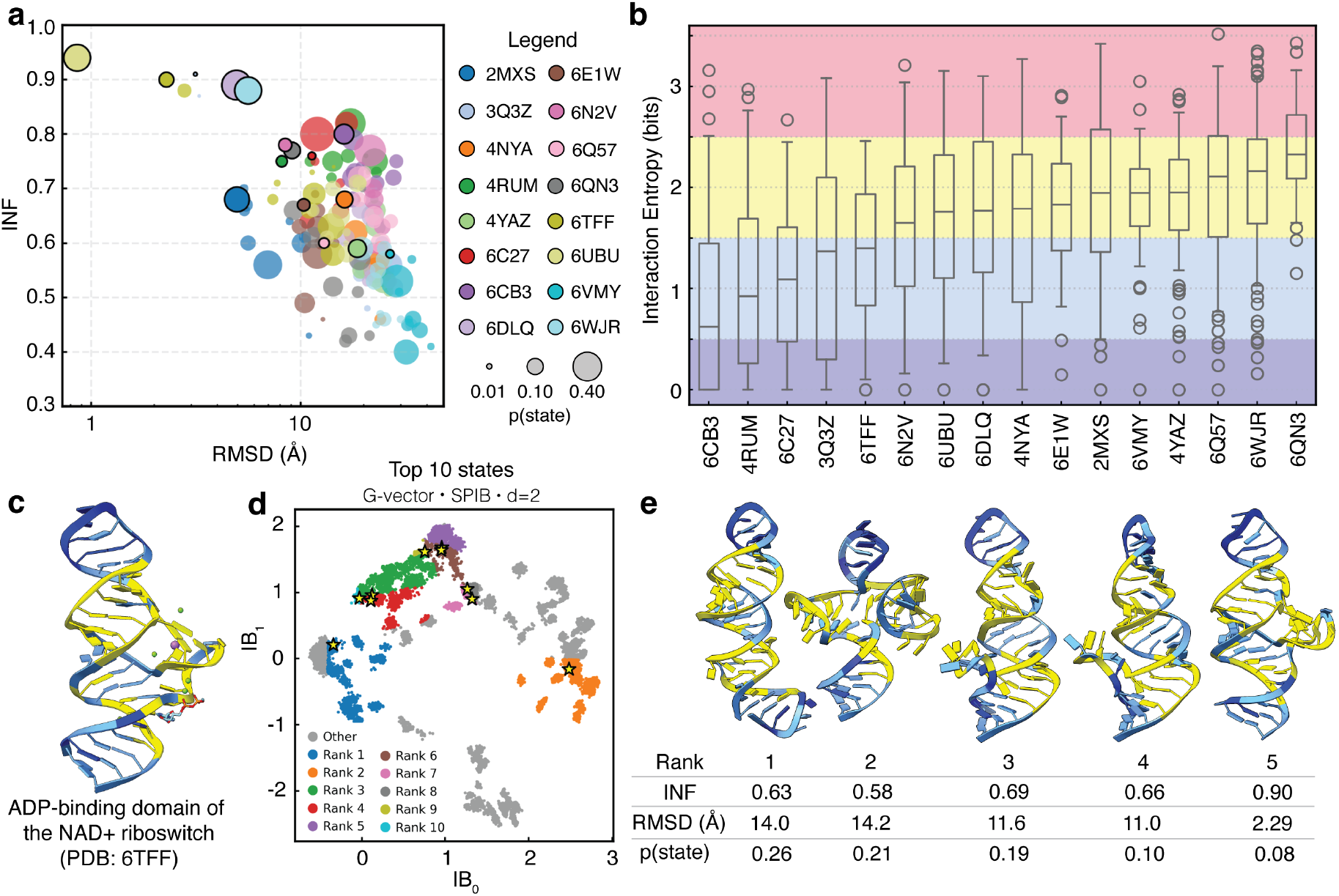
Conformational ensemble prediction with RNAnneal. **a**. Ten-state conformational ensembles are shown for each riboswitch. The location of the circles conveys the INF and RMSD of the representative conformer for the state, while the size of the circles conveys the state population. Points circled black indicate the representative with the minimum Deviation Index^26^ (RMSD/INF) from the ERC. **b**. Distribution of interaction entropies for each riboswitch. **c**. Reference ERC for the NAD^+^ riboswitch colored by the RNAnneal interaction entropy. **d**. State boundaries and representative conformers (stars) projected along the Information Bottleneck (IB). **e**. Five highest-ranked representatives for the NAD^+^ riboswitch; the INF and RMSD from the ERC and state population are provided for each representative.

**FIG. 4.**
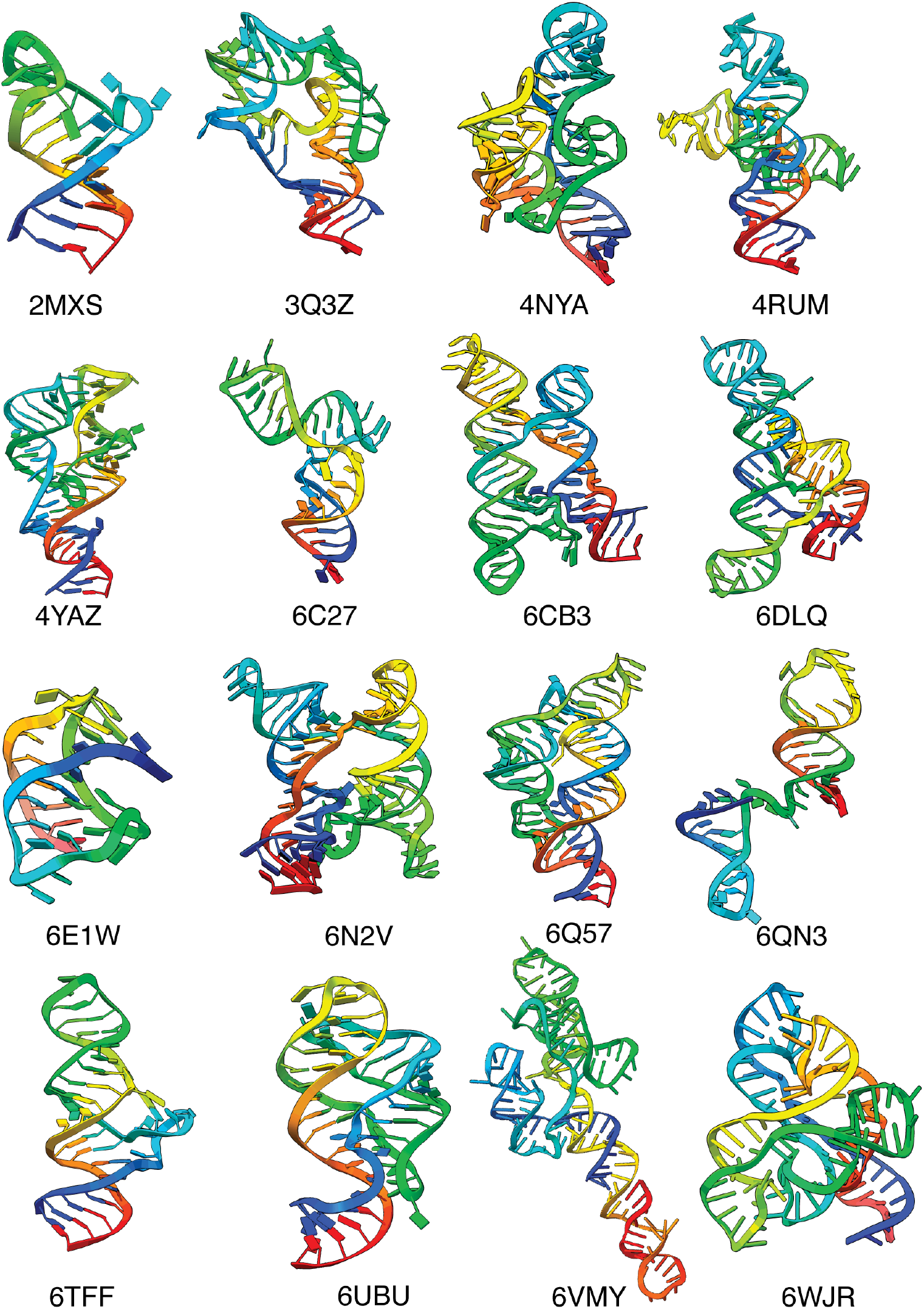
Riboswitch reference ERCs labeled with PDB accession code.

**FIG. 5.**
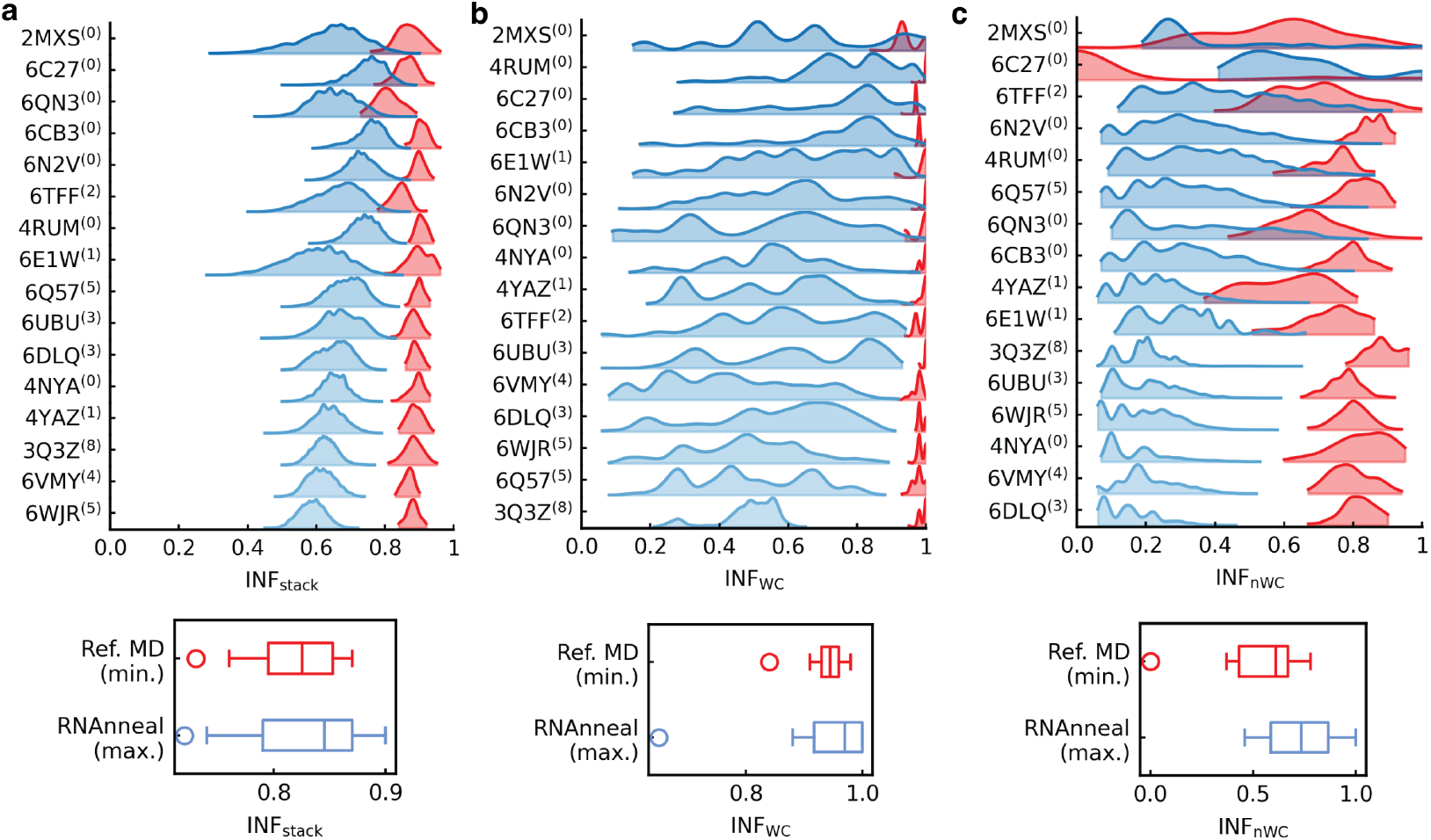
Interaction Network Fidelity (INF) by interaction type. The INF is reported for **a**. stacking interactions (Median [IQR]; Ref. MD: 0.82 [0.79 − 0.86]; RNAnneal: 0.85 [0.80 − 0.87]), **b**. Watson-Crick (WC) interactions (Ref. MD: 0.95 [0.93 − 0.96]; RNAnneal: 0.98 [0.92 − 1.0]), **c**. non-Watson-Crick (nWC) interactions (Ref. MD: 0.62 [0.48 − 0.67]; RNAnneal: 0.80 [0.59 − 0.87]). It is worth noting that the conformation with, e.g., the maximum INF_WC_, does not necessarily have the maximum INF_stack_ or INF_nWC_.

**FIG. 6.**
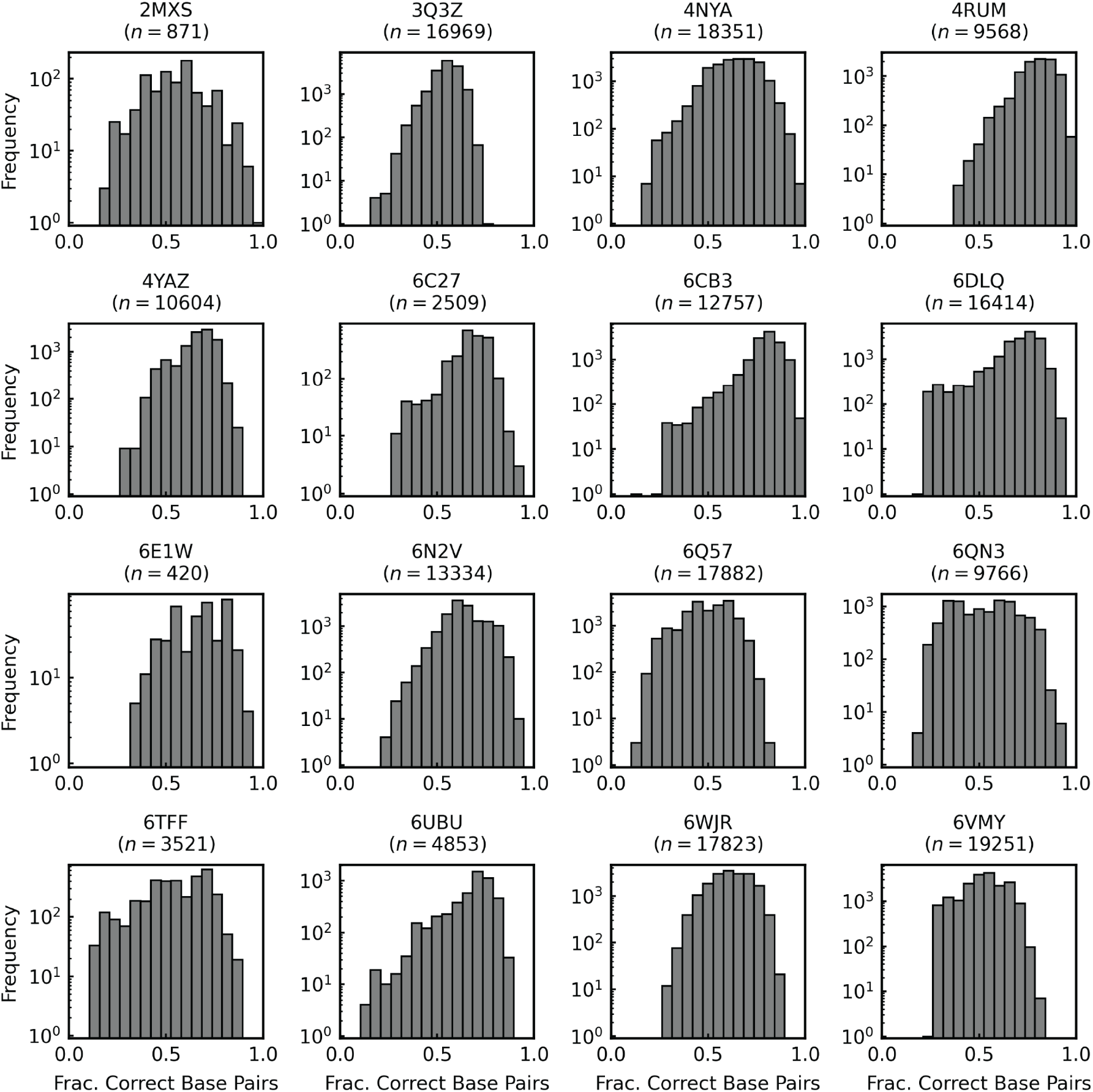
Evaluation of secondary structures predicted by EternaFold, LinearFold, and LinearAlifold. Secondary structures sampled by Eternafold, LinearFold, and LinearAilfold are compared to the reference secondary structure extracted from the listed PDB entry; *n* denotes the total number of secondary structures sampled.

**FIG. 7.**
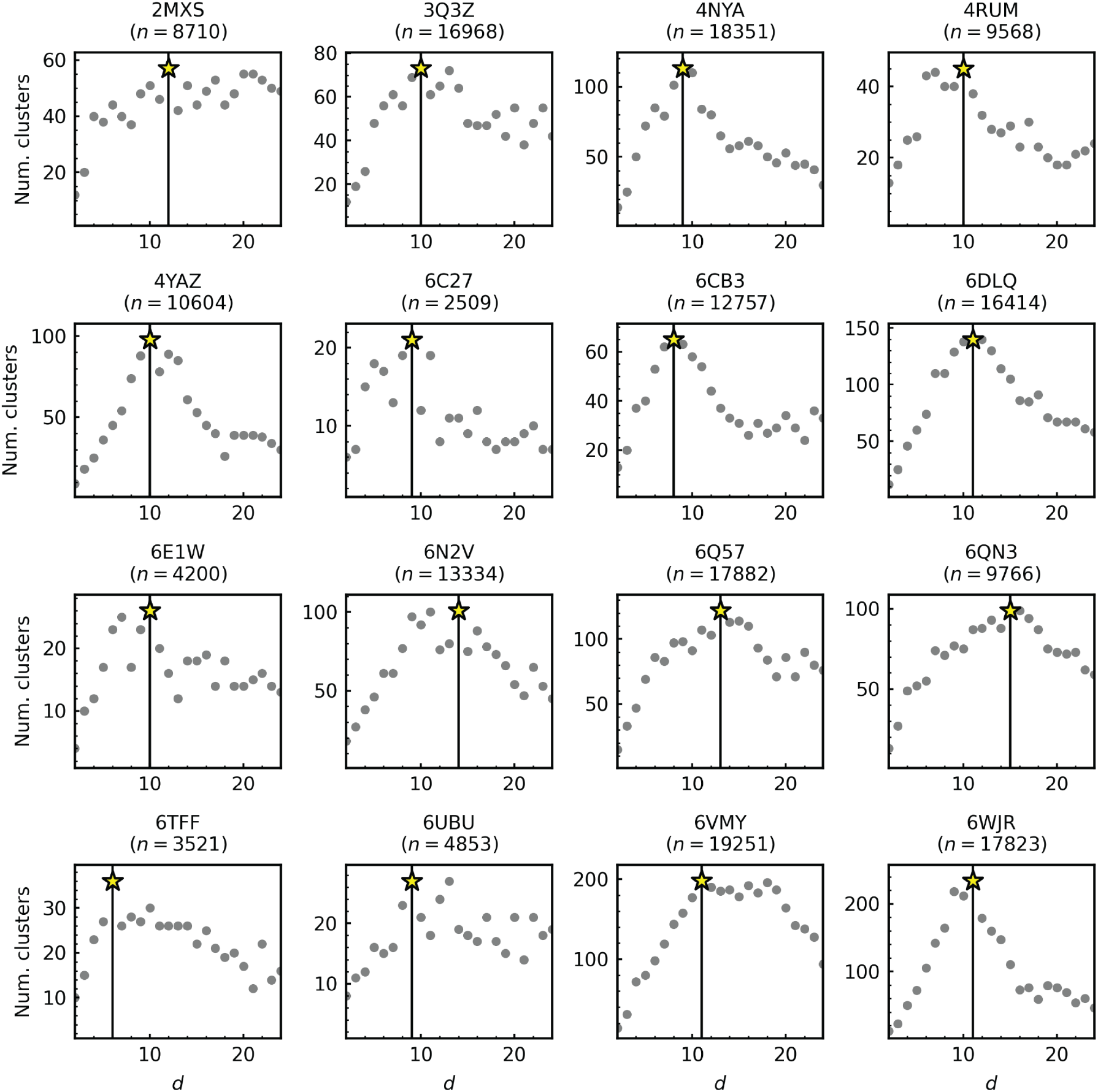
Number of clusters vs d. For each RNA, the G-vectors are projected onto *d* components, and clustering is performed via the Advanced Density Peaks algorithm implemented in dadapy. The number of clusters, *N*_*c*_ is recorded as a function of *d*. The star values and vertical line indicate the *d* where *N*_*c*_ is maximized; *n* denotes the total number of models used in the calculation.

**FIG. 8.**
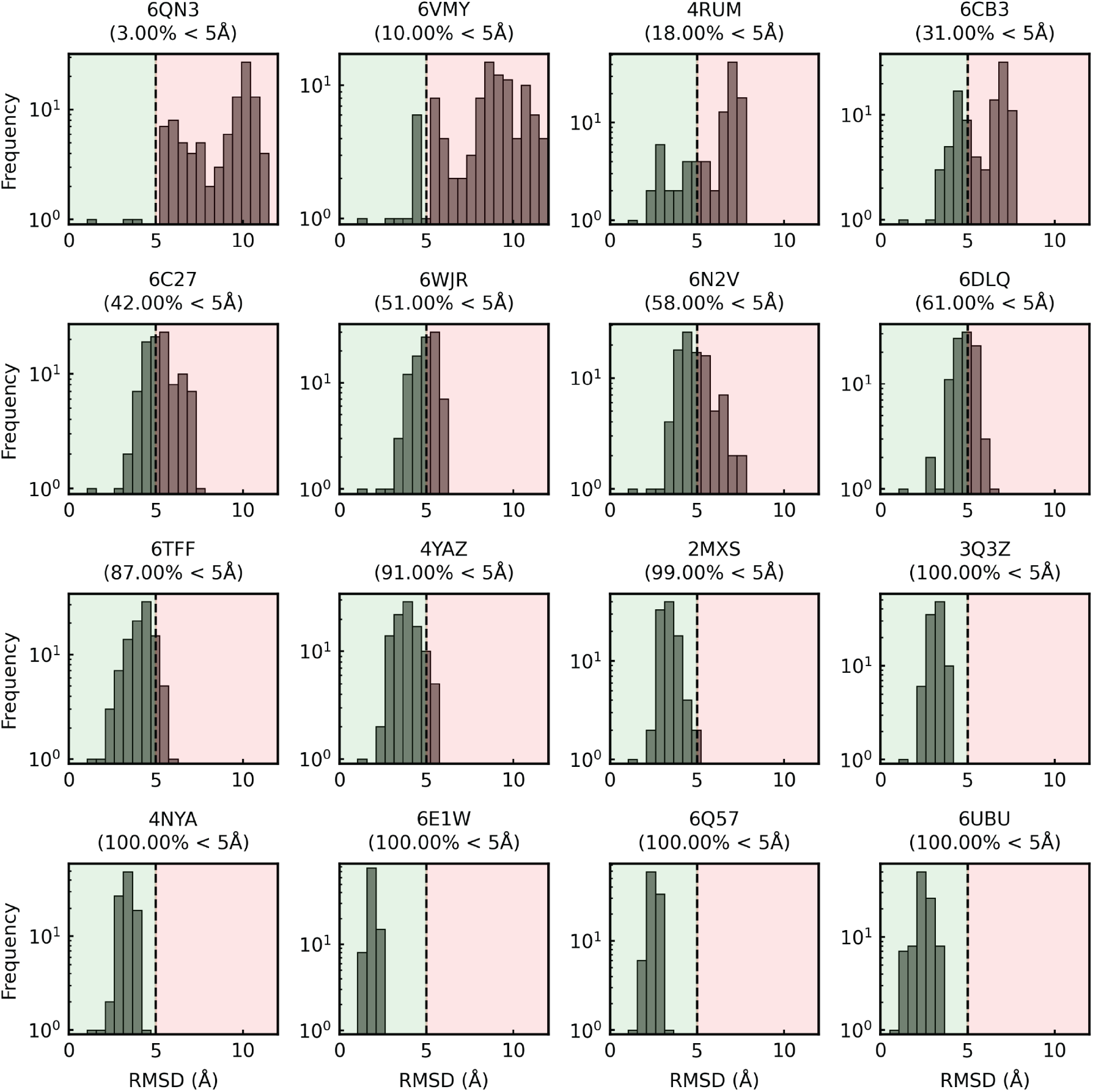
RMSD variation over a 100ns MD simulation launched from the reference conformation. The dashed line placed at 5Å indicates the cutoff used to classify ERCs (RMSD ≤ 5Å) and decoys (RMSD *>* 5Å) conformations. The proportion of ERCs is provided underneath the PDB accession code.

**FIG. 9.**
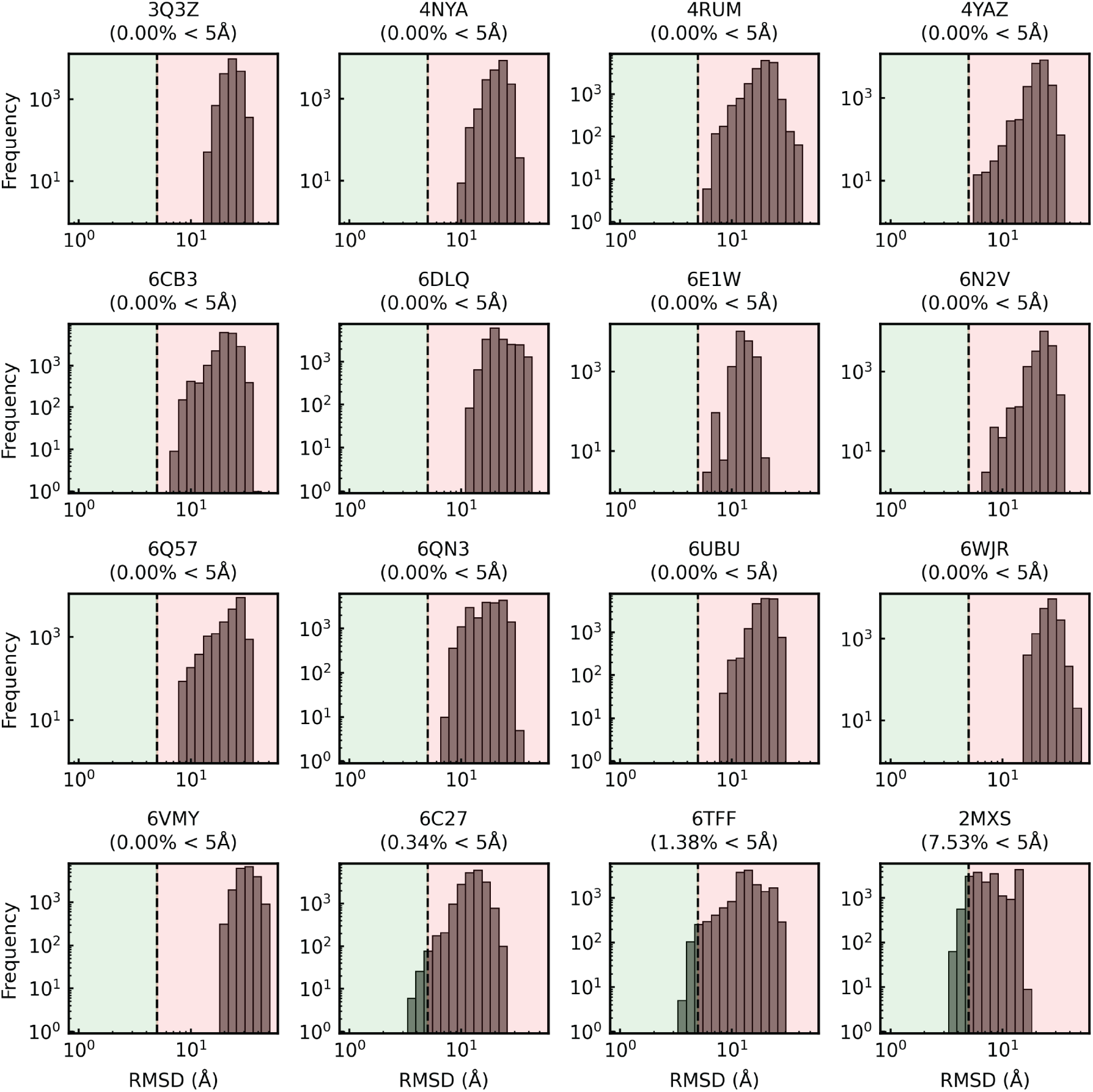
RMSD distribution of conformations sampled by RNAnneal. The dashed line placed at 5Å indicates the cutoff used to classify ERCs (RMSD ≤ 5Å) and decoys (RMSD *>* 5Å) conformations. The proportion of near-ERCs is provided underneath the PDB accession code.

**FIG. 10.**
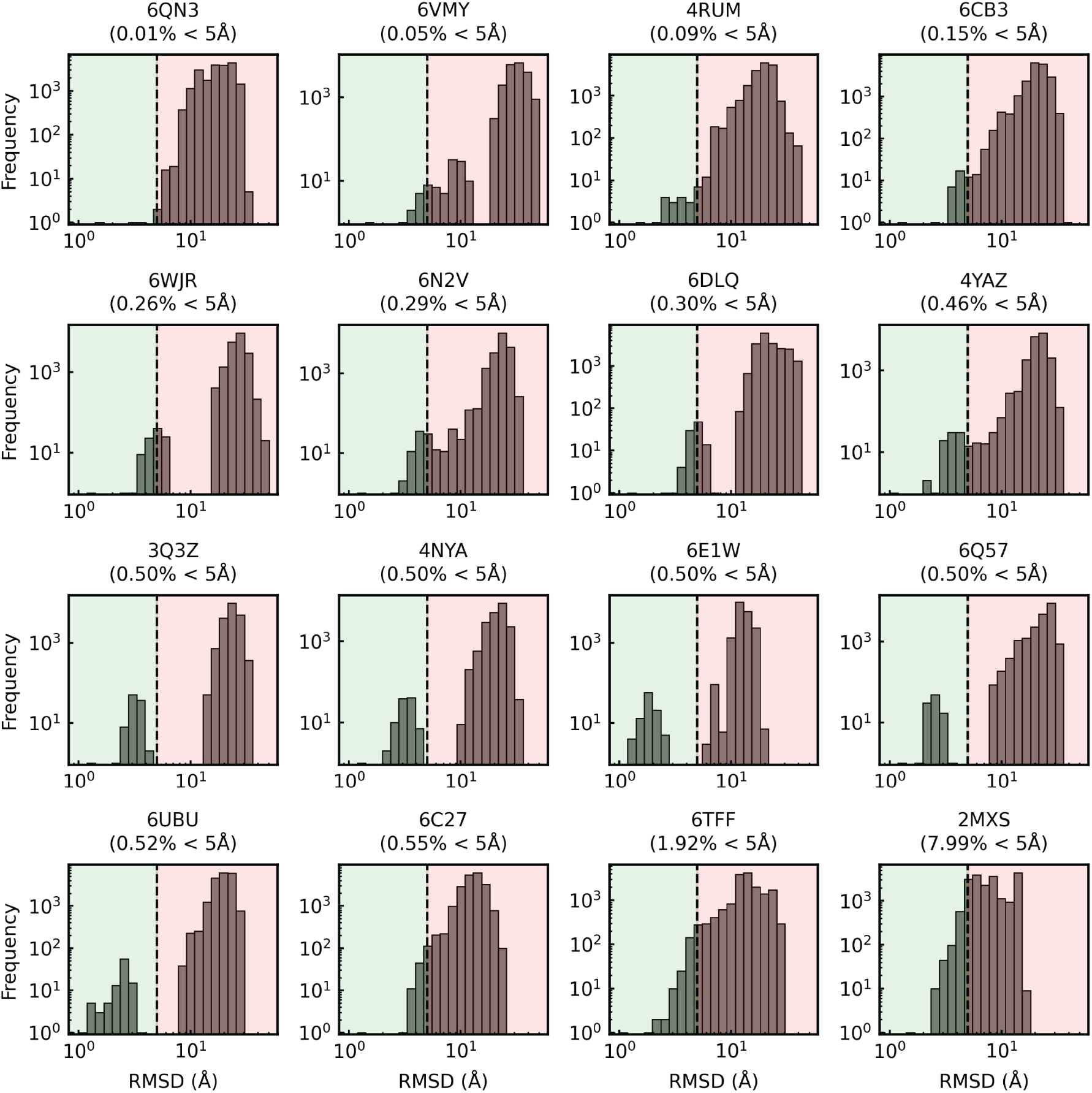
RMSD distribution of conformations for the classification task. The classification task merges structures sampled by RNAnneal with frames from simulations launched from the reference conformations (i.e., merges Fig. S8 and S9). The dashed line placed at 5Å indicates the cutoff used to classify ERC (RMSD ≤ 5Å) and decoy (RMSD*>* 5Å) conformations. The proportion of near-ERCs is provided underneath the PDB accession code.

**FIG. 11.**
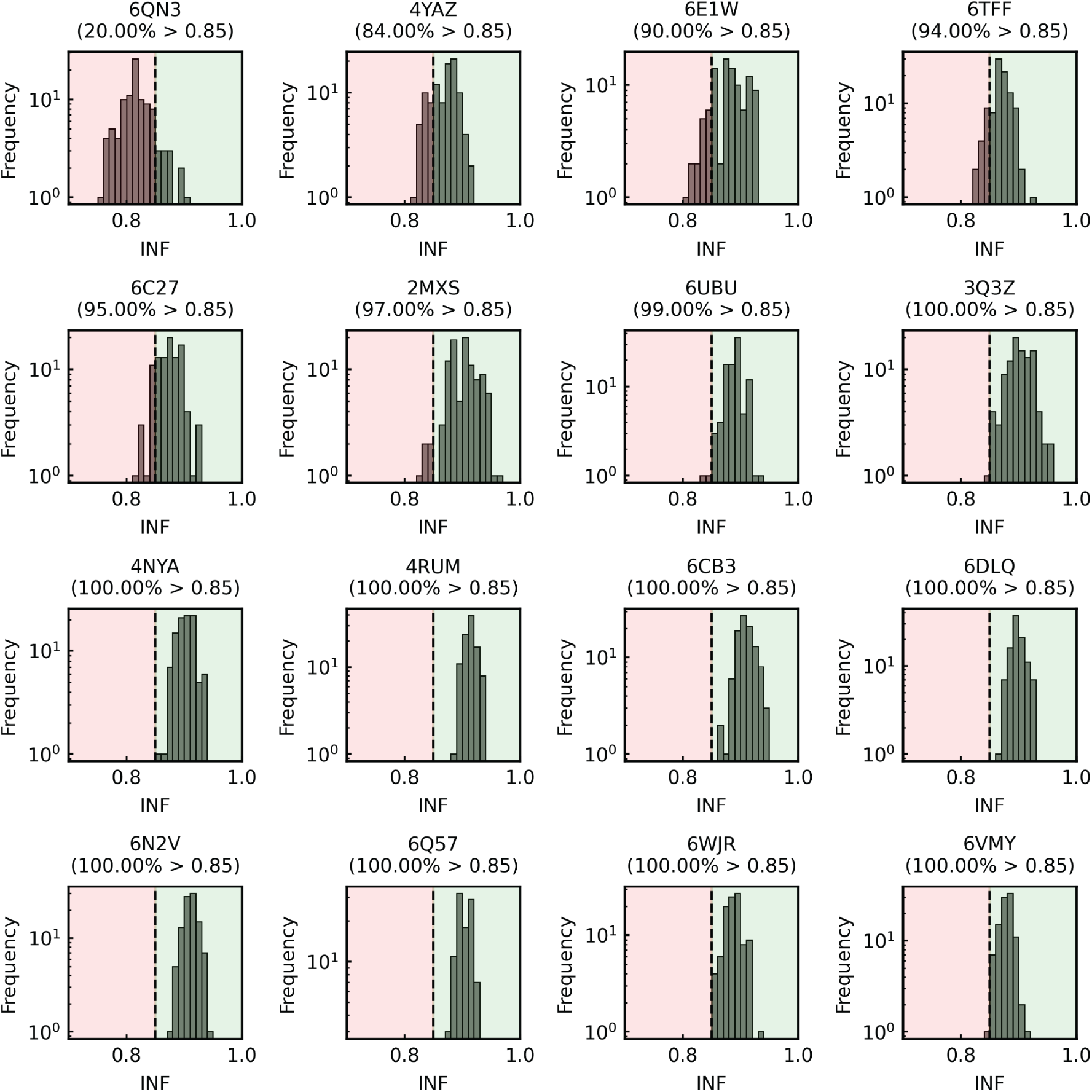
INF variation over a 100ns MD simulation launched from the reference conformation. The dashed line placed at 0.85 indicates the cutoff used to classify ERC (INF ≥ 0.85) and decoy (INF*<*0.85) conformations. The proportion of near-ERCs is provided underneath the PDB accession code.

**FIG. 12.**
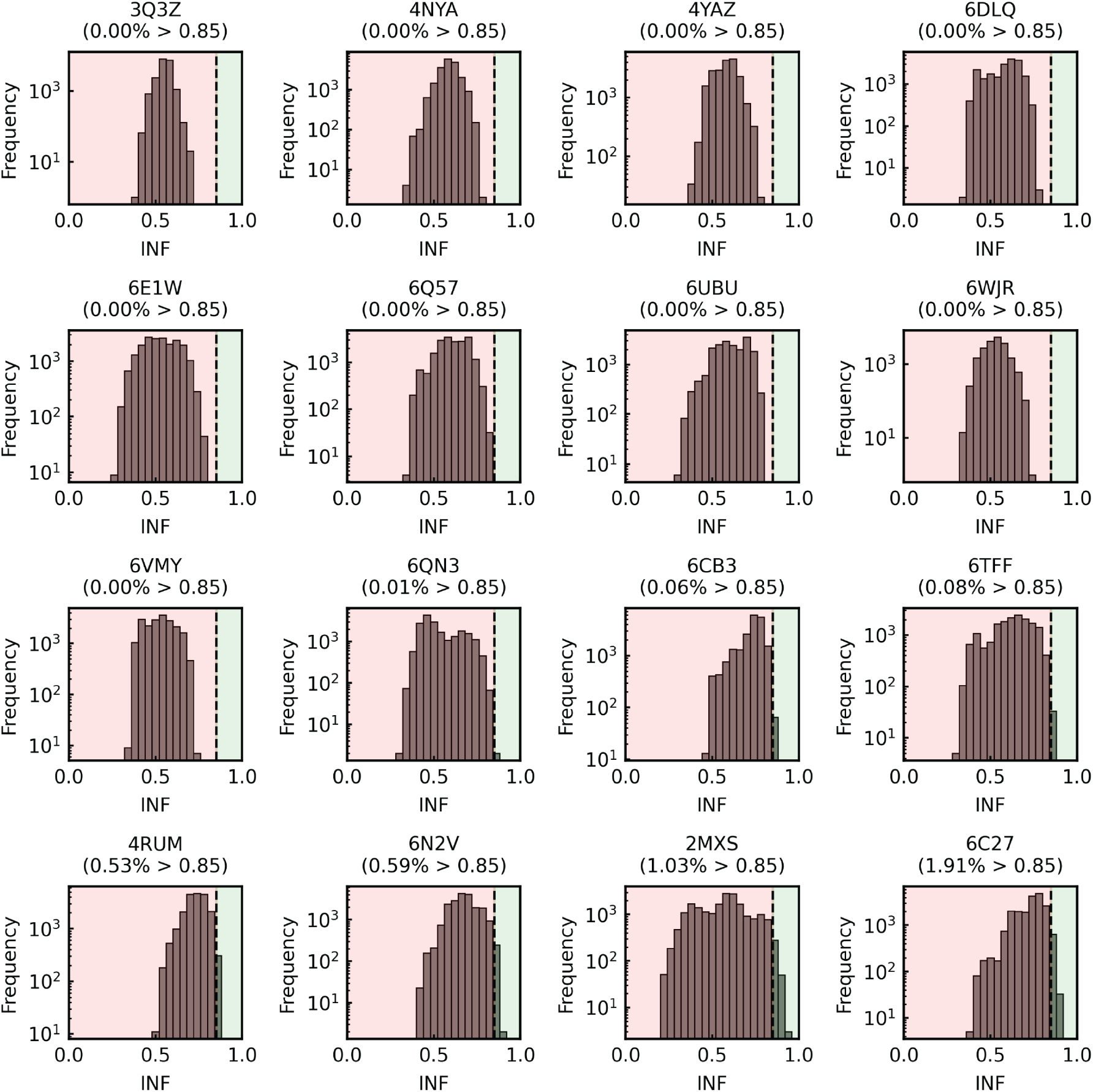
INF distribution of conformations generated by RNAnneal. The dashed line placed at 0.85 indicates the cutoff used to classify ERC (INF ≥ 0.85) and decoy (INF*<*0.85) conformations. The proportion of near-ERCs is provided underneath the PDB accession code.

**FIG. 13.**
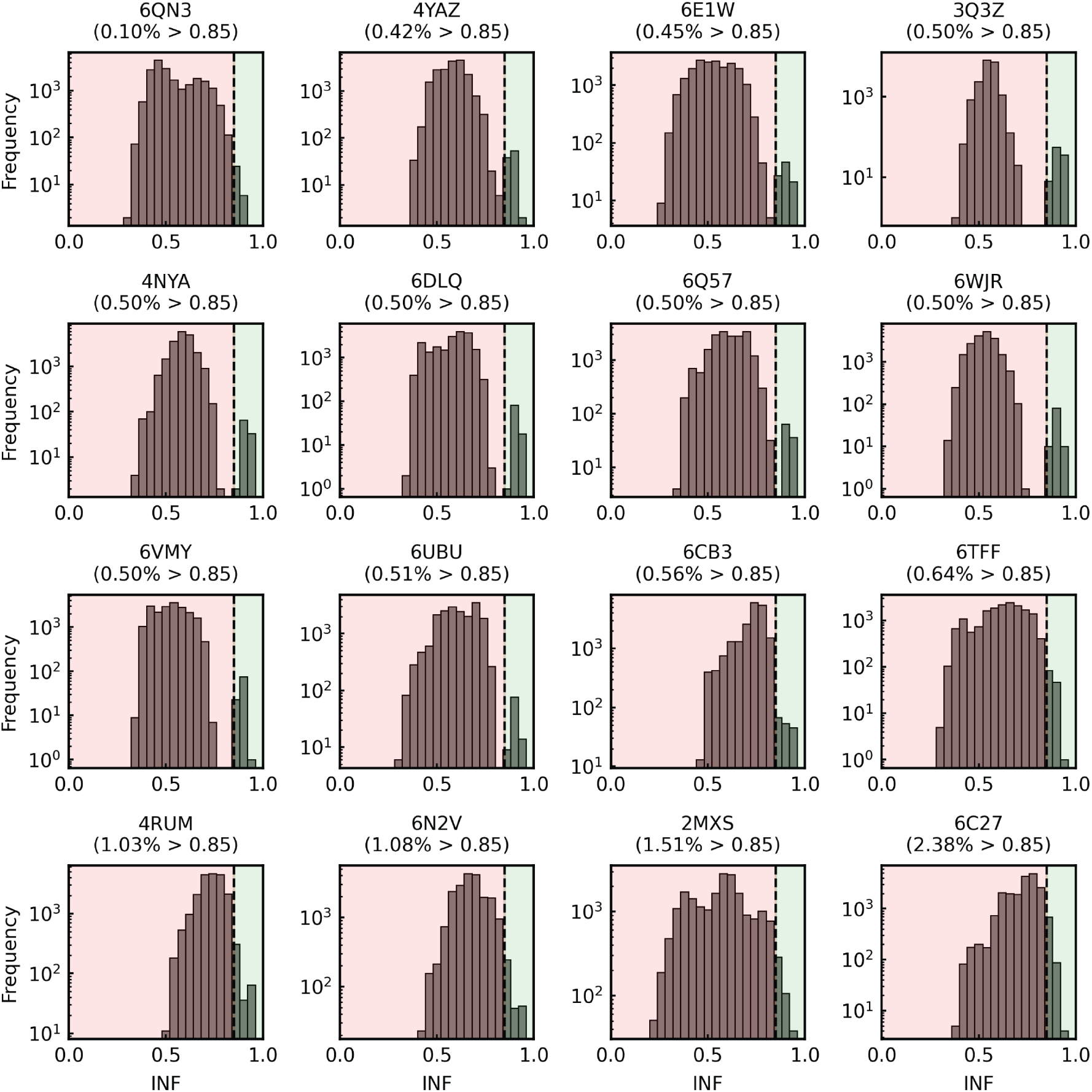
INF distribution of conformations for the classification task. The classification task merges structures sampled by RNAnneal with frames from simulations launched from the reference conformations (i.e., merges Fig. S11 and S12). The dashed line placed at 0.85 indicates the cutoff used to classify ERC (INF ≥ 0.85) and decoy (INF*<*0.85) conformations. The proportion of near-ERCs is provided underneath the PDB accession code.

**FIG. 14.**
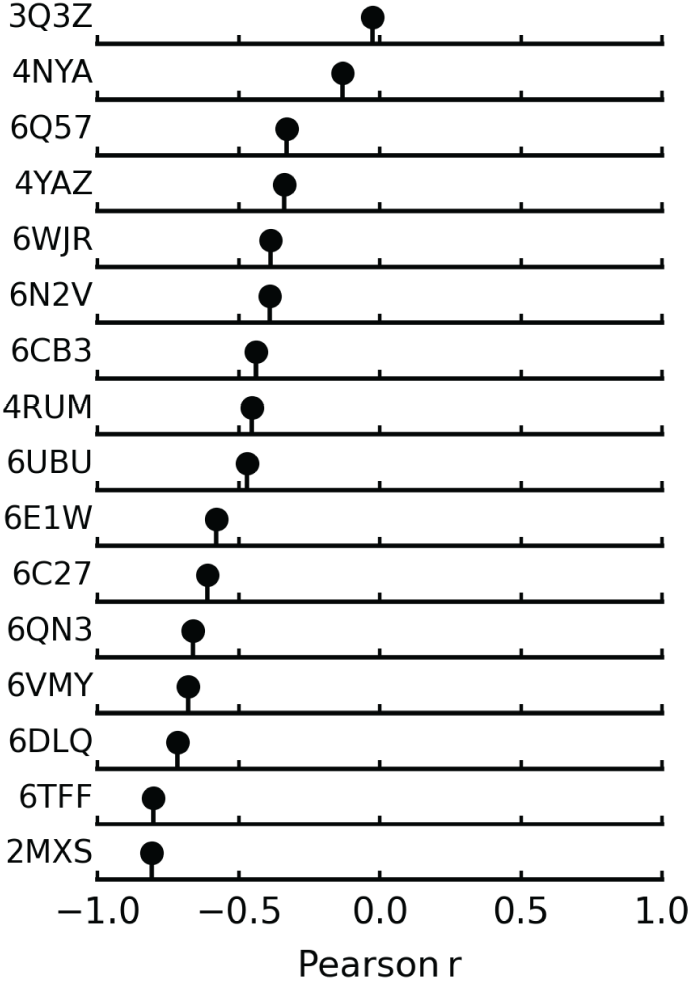
Correlation between RMSD and INF for RNAnneal conformations.

**FIG. 15.**
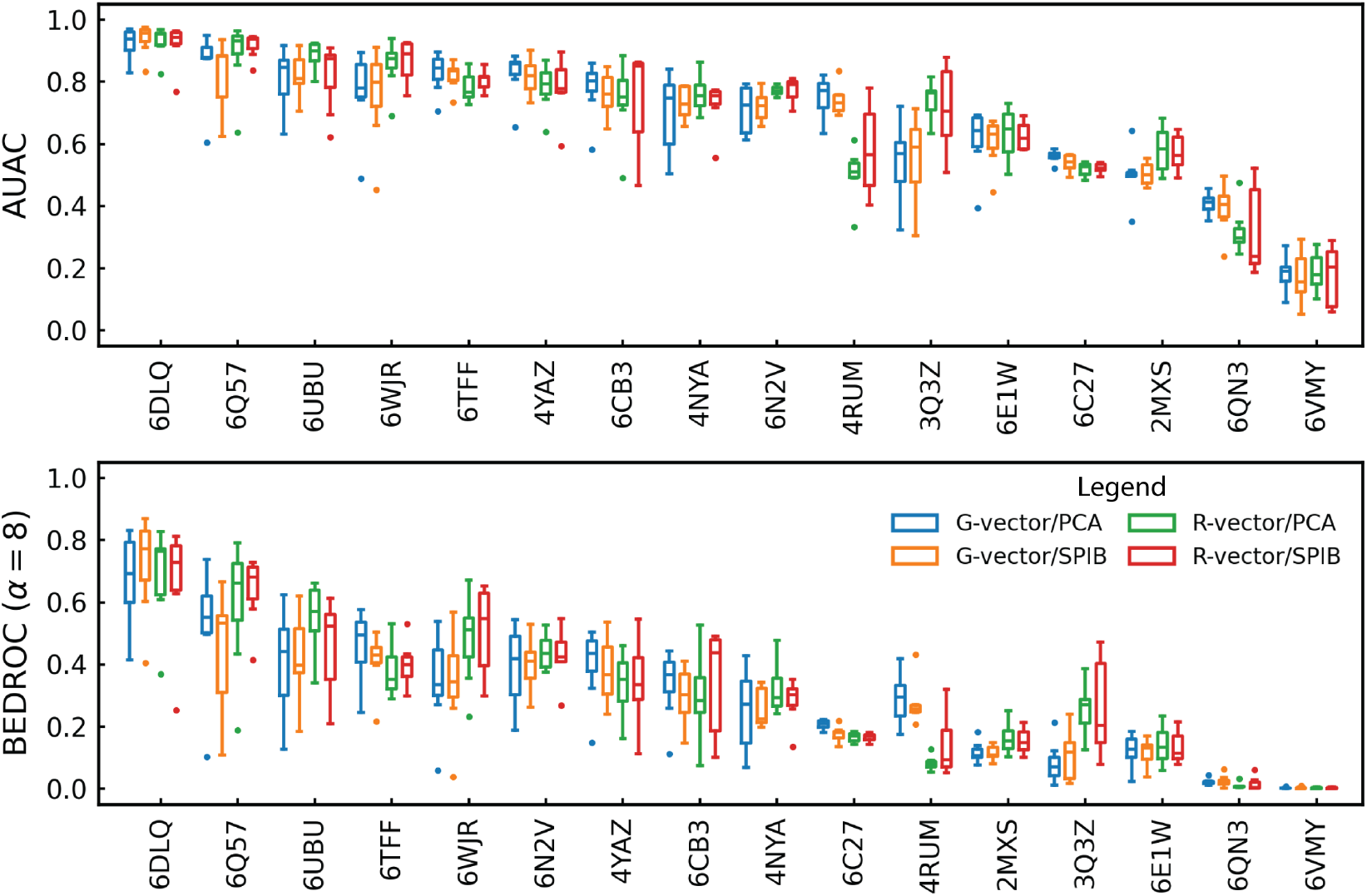
AUAC and BEDROC for each combination of training features.

**FIG. 16.**
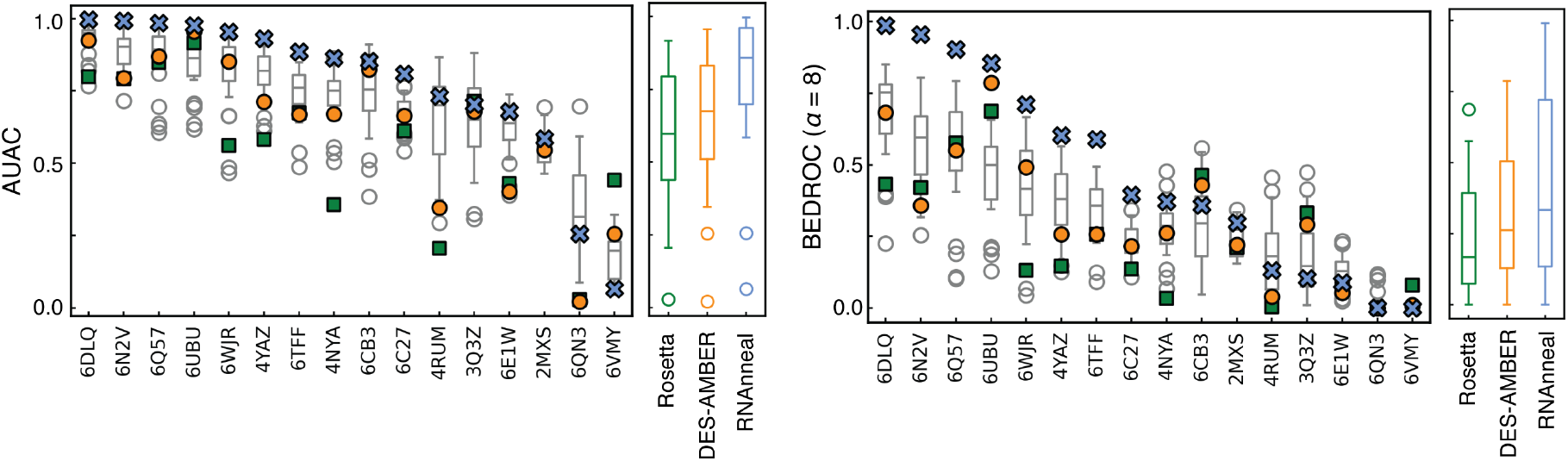
AUAC and BEDROC for RMSD classification criteria. Classification power of the Rosetta (blue), DES-AMBER (orange), and RNAnneal (blue) scores when ERCs are defined as conformations *<* 5Å in RMSD from the reference conformation. The gray boxplots show the distribution of AUAC and BEDROC values for each model in the TM ensemble.

## Unsupervised sampling and prediction of RNA structures

RNAnneal predicts RNA conformations by sequentially sampling and scoring candidate structure models. As such, the first step is to assemble a diverse, physically plausible candidate set of 3D models (Fig. 1a). With only the primary sequence as input, RNAnneal employs physics-based and bioinformatics programs to generate a large number of secondary structures (Eternafold^16^, Linearfold^17^ and LinearAlifold^18^; see Secondary Structure Prediction), which are then assembled into all-atom 3D models using the FARFAR2 algorithm (see FAR-FAR2).^16–19^ Next, 200 diverse, representative models of the 3D structure are selected by clustering using geometric features and ranking conformations based on their Rosetta scores (see Clustering). To produce a final set of candidate structures, each of the representative conformations is simulated for 100ns via implicit solvent MD (see Molecular Dynamics).

Next, we describe the RNAnneal score, which is extracted from an ensemble of generative models trained on energy values (see Molecular Dynamics) and latent representations of candidate conformations (Fig. 1b). To arrive at the latent representations, geometric features (see R- and G-vectors) are extracted from the candidate conformations and subsequently encoded as latent representations by Principal Component Analysis (PCA) and a State Predictive Information Bottleneck^20^ (see SPIB) model; the principal components capture the highest-variance geometric features, while the SPIB projections preserve the slowest processes of the MD simulations and learn state boundaries that are metastable over a user-specified timescale.

Then, an ensemble of Thermodynamic Maps^12^ (TMs; see Thermodynamic Maps) is trained to estimate the log-likelihood of the candidate conformations based on their energies and latent representations (Fig. 1c).^12^ In this work, we train 28 TMs on combinations of features (R- and G-vectors) and projections (SPIB and PCA) with 2 to 8 latent dimensions, *d* (Fig. 1d). It is worth noting that the energy values are consistent across all models, even though the latent representations of a given conformation vary from model to model. Finally, for a given conformation, the TM log-likelihoods are averaged to produce the RNAnneal score (see RNAnneal Score). Our evaluations found that the RNAnneal score can reliably distinguish between ERCs and decoys (Fig. 1e and Fig. 2c-d).

Since the TM models represent the energy scale of the conformational landscape as a temperature, one may anneal the RNAnneal score by adjusting the temperature of the prior distribution of the TMs during the loglikelihood calculation (see RNAnneal Score). We investigated the classification power of the RNAnneal score as a function of temperature and found that ERCs are most prioritized at low temperatures, which is consistent with ERCs lying low on the free energy landscape (Fig. 17).

**FIG. 17.**
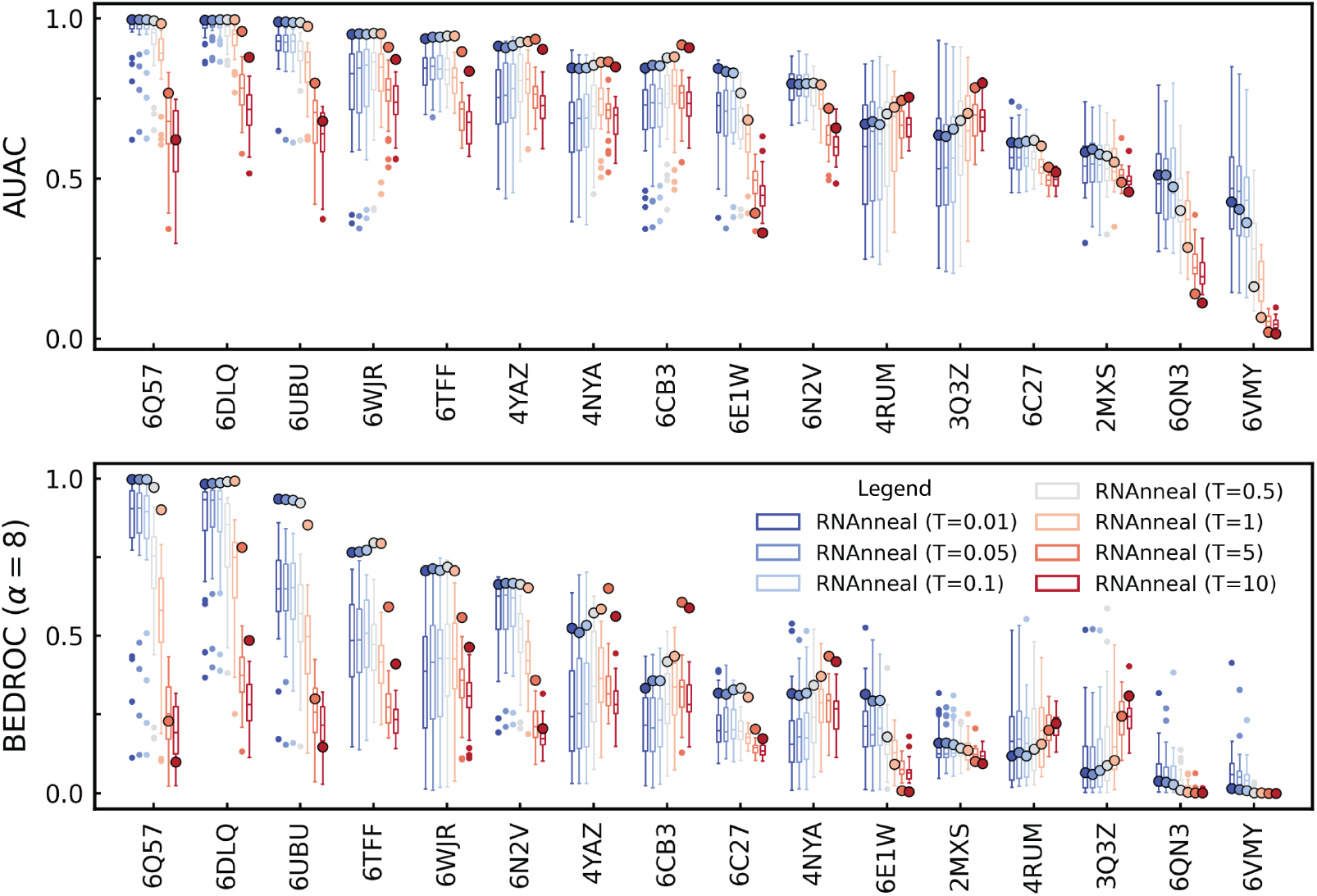
Variation in AUAC and BEDROC with scoring temperature.

**FIG. 18.**
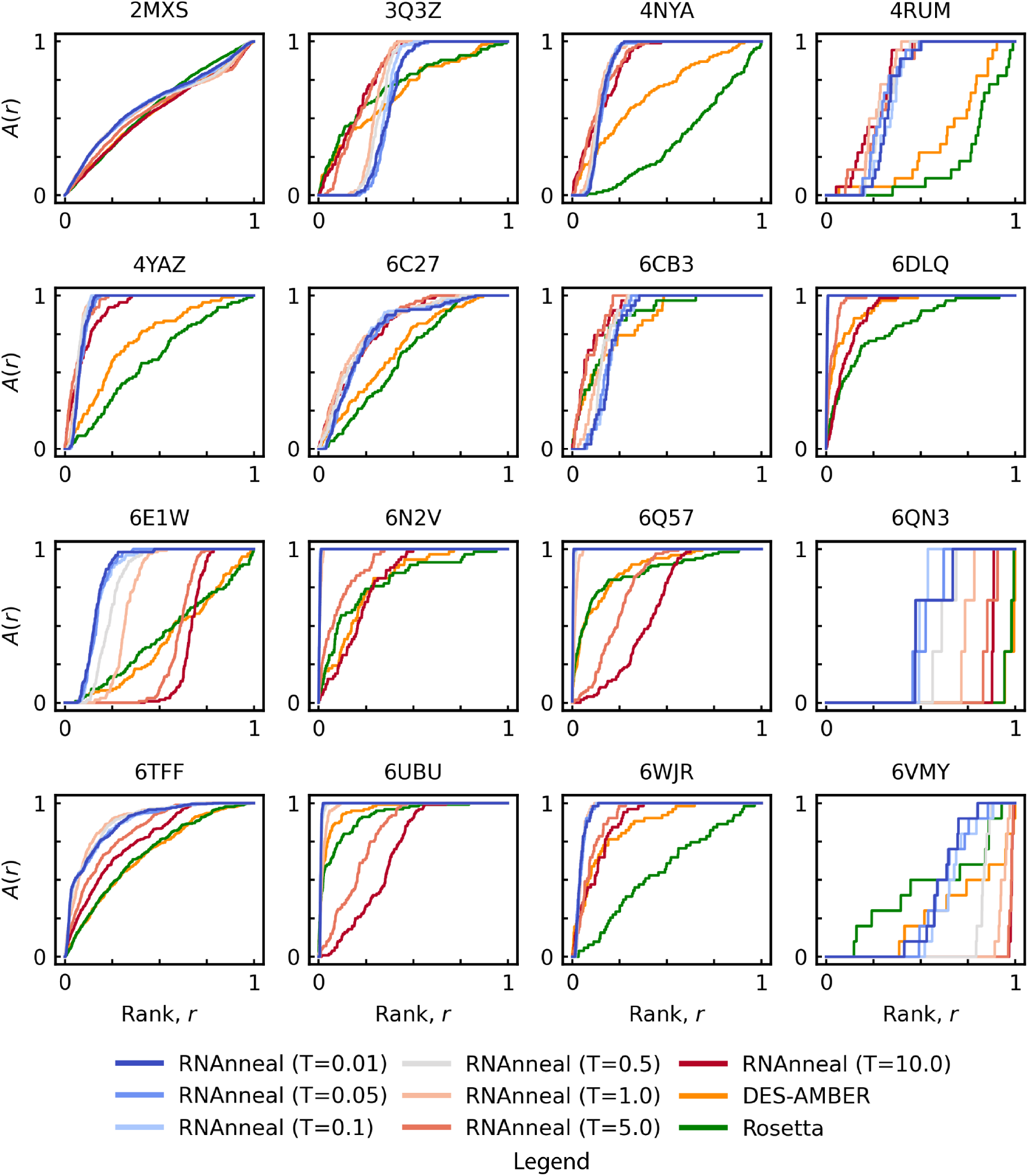
Accumulation Curves. Accumulation curves, *A*(*r*) are reported for each riboswitch, identified by the PDB entry of the reference conformation.

**FIG. 19.**
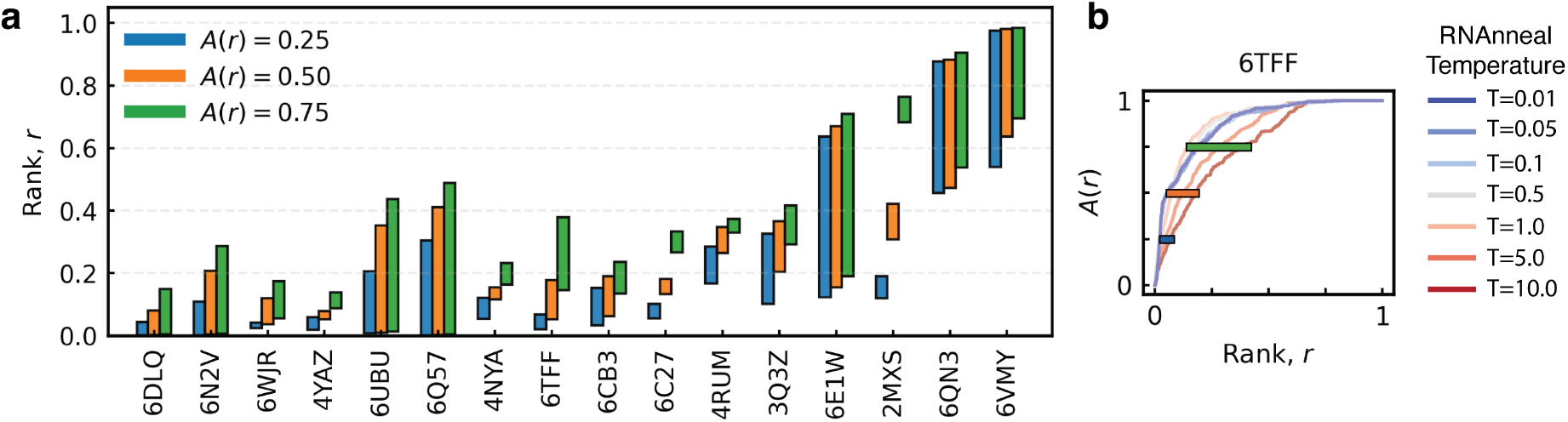
Summary of Accumulation Curves a. The set of accumulation curves for each riboswitch PDB is shown with the range of rank, *r*, for which *A*(*r*) = 0.25, *A*(*r*) = 0.50, and *A*(*r*) = 0.75. b. The range indicated by the bars shows the variation of *r* with temperature at fixed values of *A*(*r*) = 0.25, *A*(*r*) = 0.50, and *A*(*r*) = 0.75. The accumulation curve for the NAD^+^ riboswitch (6E1W^33^) is shown as an example (see Fig. S18 for all riboswitches).

**FIG. 20.**
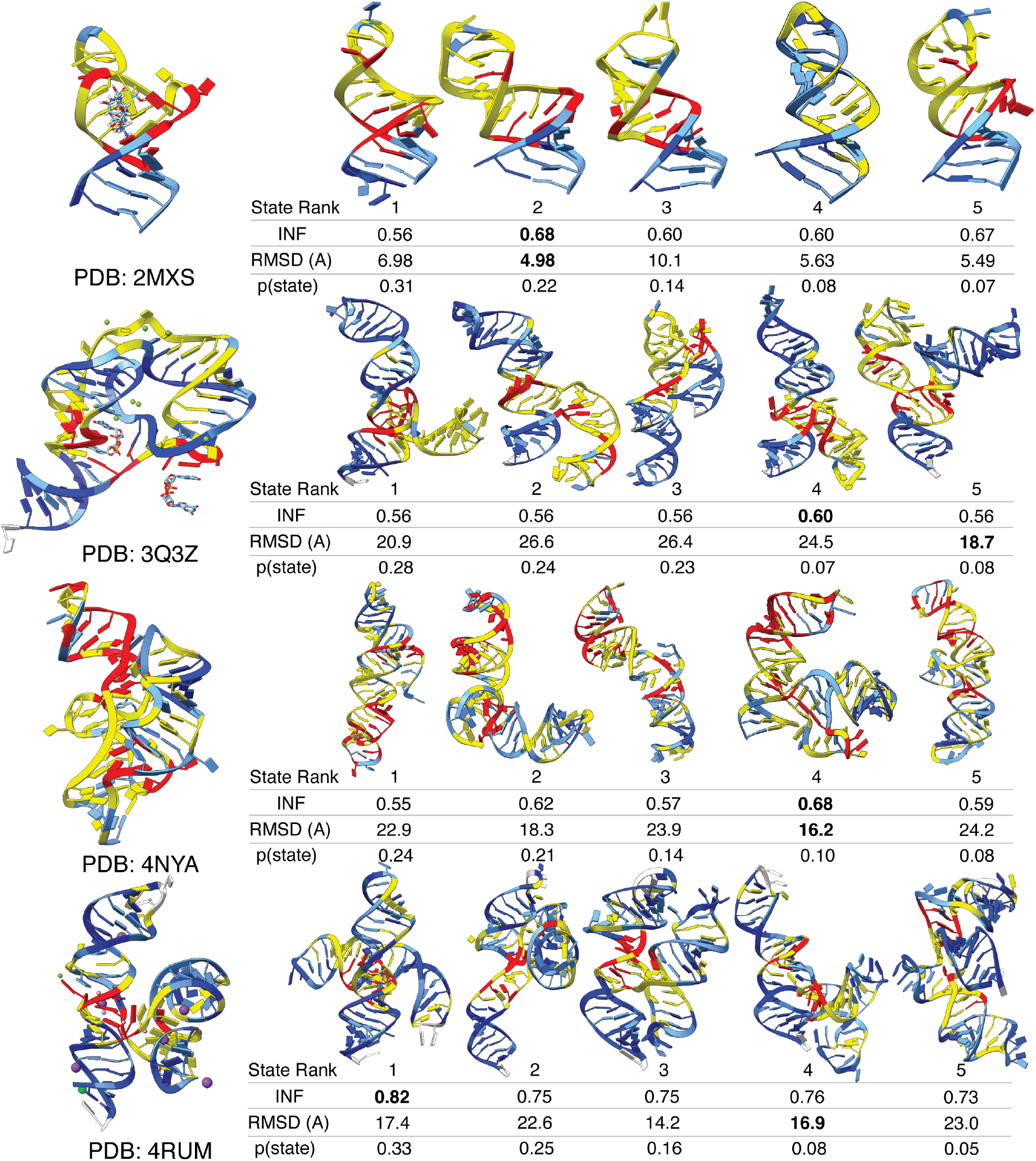
Conformational ensembles predicted by RNAnneal (2MXS, 3Q3Z^43^, 4NYA^41^, 4RUM^30^). The reference ERC for each RNA are depicted to the left. The five most probable states are reported alongside their probabilities. The best scoring model from each state is shown as a representative conformation, and the INF and RMSD values compared to the ERC are reported. Nucleotides are colored based on the interaction entropy (IE) predicted by RNAnneal according to the ranges marked in Fig. 3b (dark blue: 0 ≤ IE *<* 0.5; light blue: 0.5 ≤ IE *<* 1.5; yellow: 1.5 ≤ IE *<* 2.5; red: IE ≥ 2.5)

**FIG. 21.**
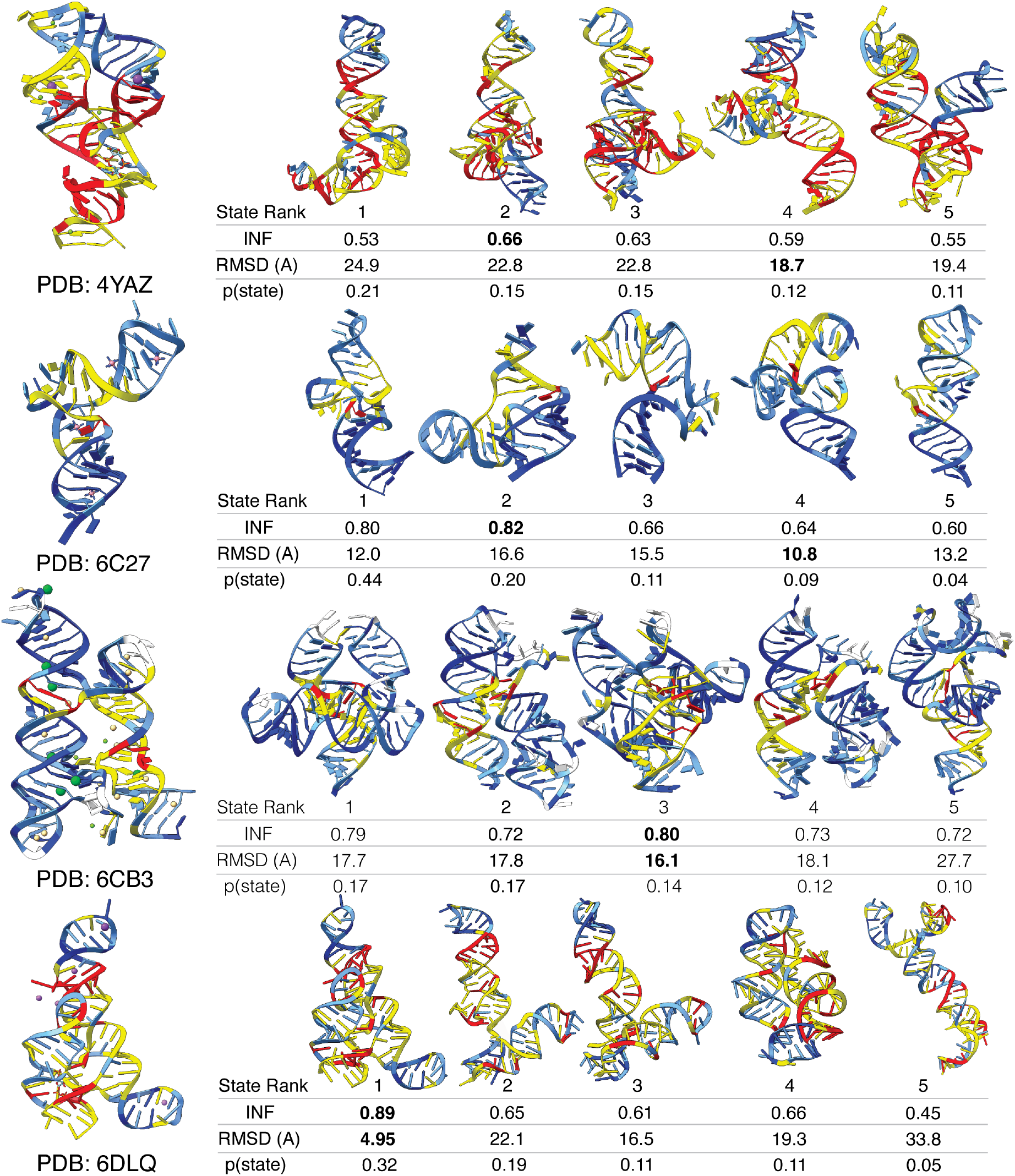
Conformational ensembles predicted by RNAnneal (4YAZ^42^, 6C27^39^, 6CB3^31^, 6DLQ^35^). The reference ERC for each RNA are depicted to the left. The five most probable states are reported alongside their probabilities. The best scoring model from each state is shown as a representative conformation, and the INF and RMSD values compared to the ERC are reported. Nucleotides are colored based on the interaction entropy (IE) predicted by RNAnneal according to the ranges marked in Fig. 3b (dark blue: 0 ≤ IE *<* 0.5; light blue: 0.5 ≤ IE *<* 1.5; yellow: 1.5 ≤ IE *<* 2.5; red: IE ≥ 2.5)

**FIG. 22.**
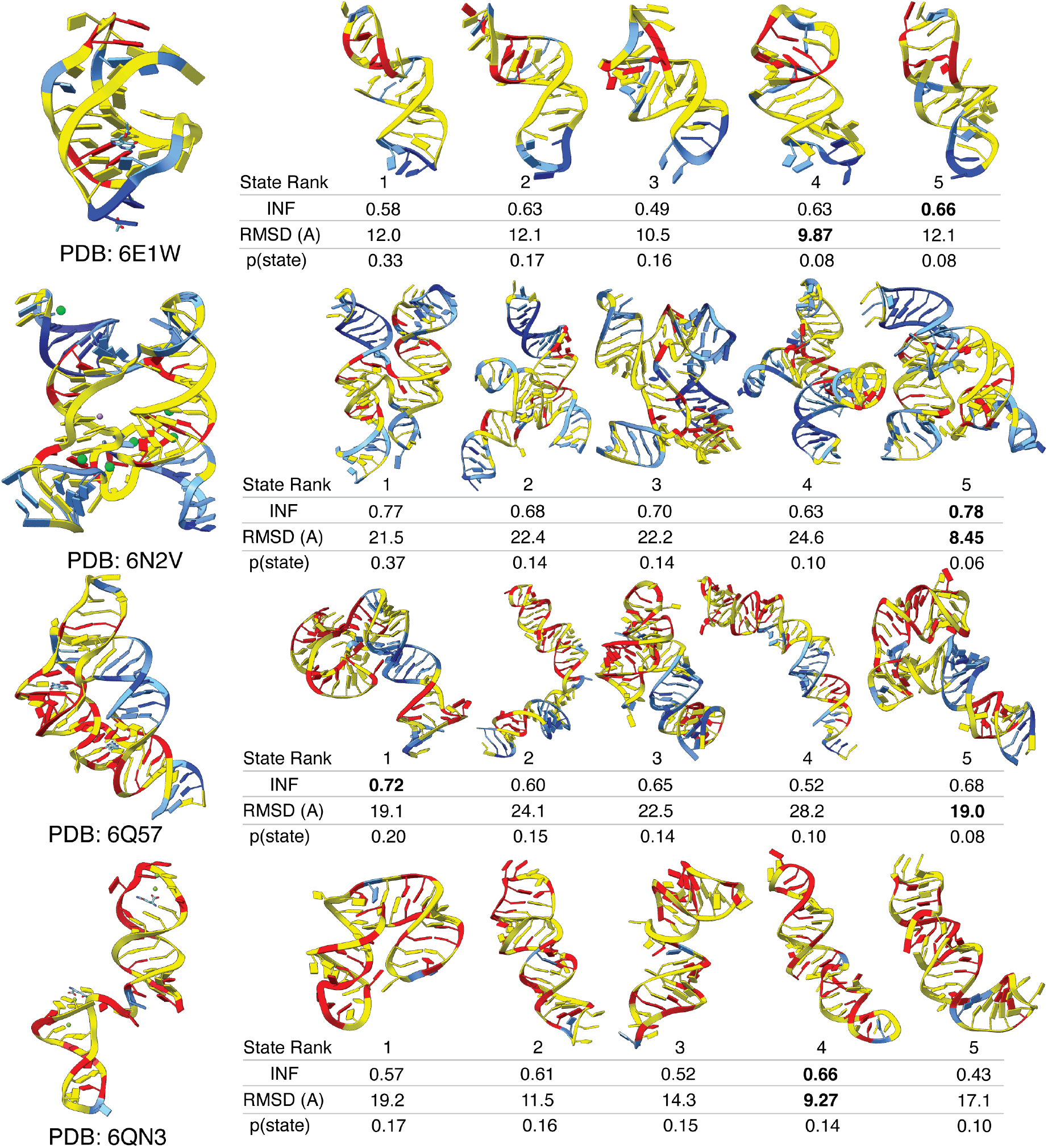
Conformational ensembles predicted by RNAnneal (6E1W^33^, 6N2V^29^, 6Q57^36^, 6QN3^32^). The reference ERC for each RNA are depicted to the left. The five most probable states are reported alongside their probabilities. The best scoring model from each state is shown as a representative conformation, and the INF and RMSD values compared to the ERC are reported. Nucleotides are colored based on the interaction entropy (IE) predicted by RNAnneal according to the ranges marked in Fig. 3b (dark blue: 0 ≤ IE *<* 0.5; light blue: 0.5 ≤ IE *<* 1.5; yellow: 1.5 ≤ IE *<* 2.5; red: IE ≥ 2.5)

**FIG. 23.**
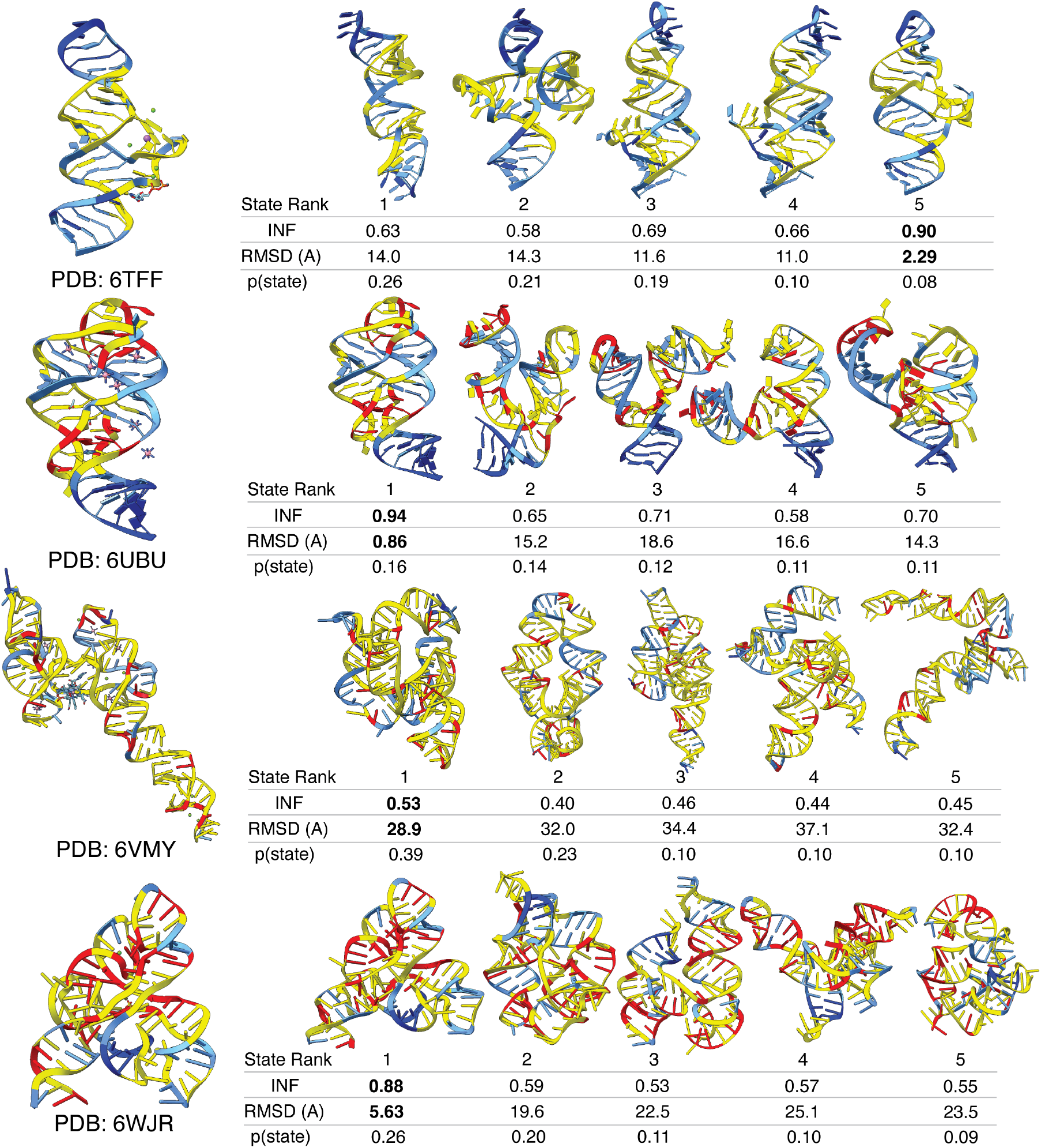
Conformational ensembles predicted by RNAnneal (6TFF^27^, 6UBU^37^, 6WJR^38^, and 6VMY^40^). The reference ERC for each RNA are depicted to the left. The five most probable states are reported alongside their probabilities. The best scoring model from each state is shown as a representative conformation, and the INF and RMSD values compared to the ERC are reported. Nucleotides are colored based on the interaction entropy (IE) predicted by RNAnneal according to the ranges marked in Fig. 3b (dark blue: 0 ≤ IE *<* 0.5; light blue: 0.5 ≤ IE *<* 1.5; yellow: 1.5 ≤ IE *<* 2.5; red: IE ≥ 2.5)

## Results

RNAnneal’s structure prediction performance is evaluated on a set of 16 experimentally determined riboswitch structures originally curated in Ref.^21^ (see Reference Structures; Table I and Fig. S4). Riboswitches are typically found in the 5’-untranslated region of bacterial mRNAs and are *cis*-acting regulatory elements that control gene expression by up- or down-regulating transcription or translation.^22^ The basis of riboswitch function is conformational: aptamer domains fold into structurally distinct “bound” (*holo*) and “free” (*apo*) states in the presence or absence of a cognate ligand, which in turn transmits a signal that alters expression. Riboswitches bind their cognate ligands with high specificity through higher order structural motifs, including multi-helix junctions, loop-loop interactions, and pseudoknots. Such motifs are rich in non-WC interactions and their structures are typically challenging to model prospectively.^23–25^

**TABLE 1.**
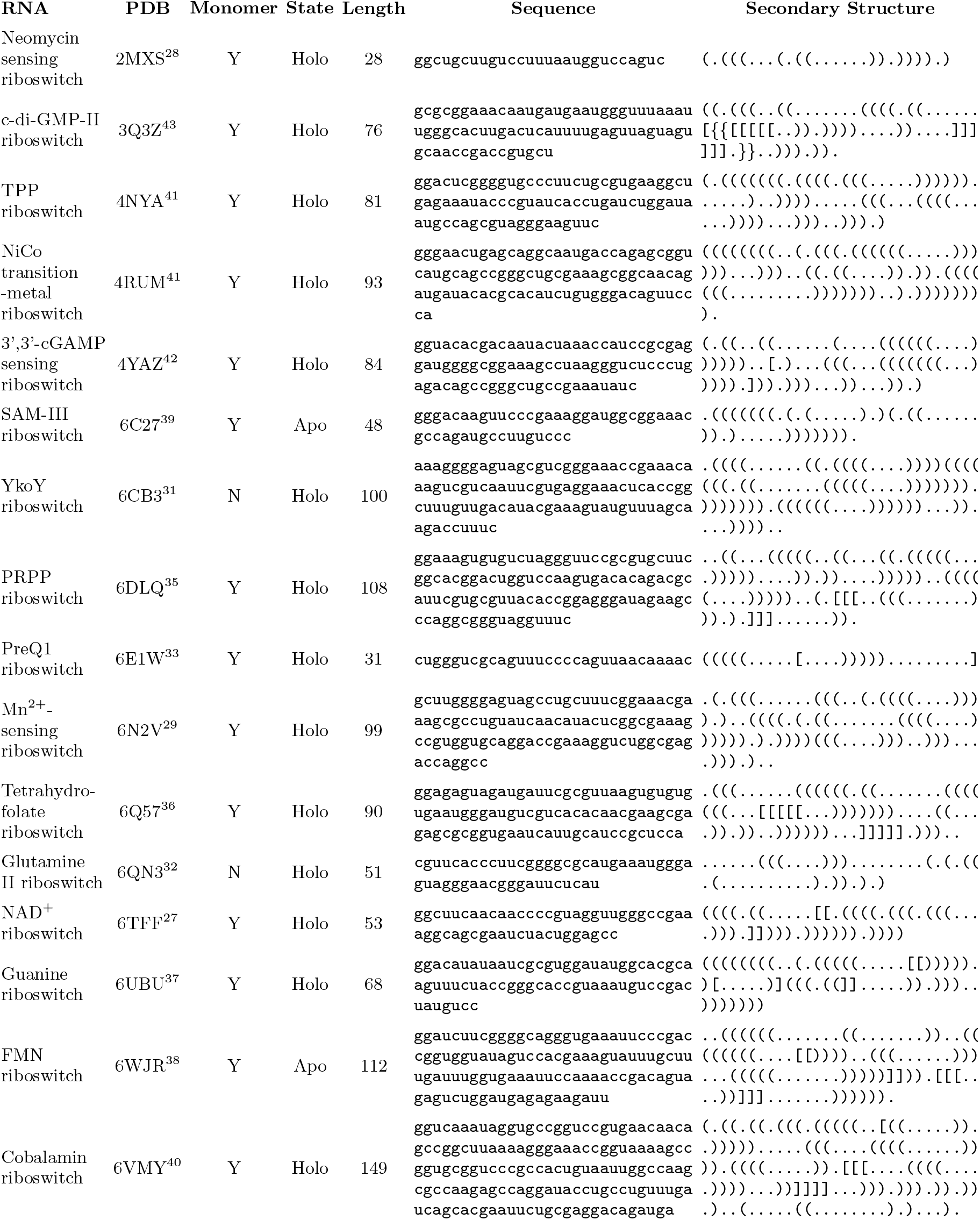
Details of riboswitch sequences and reference conformations.

The Root Mean Squared Deviation (see RMSD) and Interaction Network Fidelity (see INF) are employed to measure structural discrepancy between RNAnneal conformations and ERCs. The RMSD measures the average change in atomic positions after optimal alignment of two conformations, while the INF measures the correlation between their nucleobase interaction networks. Both metrics are required because minor differences in the interaction network (e.g., at a helix junction) may result in large coordinate changes and therefore a large RMSD. The INF is more robust to such variations since no alignment is required.

To summarize, we found that RNAnneal consistently samples PK-free near-ERCs, and its score prioritizes them over decoys. Next, we coarse-grained the RNA structures predicted by RNAnneal into states with the goal of predicting weighted ensembles of representative conformations. Then, to assess conformational heterogeneity, we introduced the interaction entropy to quantify the variability of a given nucleotide’s interaction patterns within an ensemble. Finally, we show that annealing the RNAnneal score improved ERC classification for most of the riboswitches

### RNAnneal samples experimentally resolved conformations of pseudoknot-free riboswitches

First, we assess if RNAnneal’s conformational sampling procedure (Fig. 1a) produces conformations that are similar to ERCs. We compared the interaction composition of conformations sampled by RNAnneal to the reference ERCs and found that 60-80% of interactions are stacking, 10-30% are Watson-Crick (WC), and 0-20% are non-WC for both sets (Fig. 2a). The interaction composition of the ERCs is consistent, even though their lengths vary from 28 to 149 nucleotides (Fig. 2b).

Next, the structural similarity of RNAnneal conformations to the ERC was evaluated using RMSD and INF. To assess the stability of the ERCs with respect to thermal fluctuations, we simulated each one for 100ns (see Molecular Dynamics) and monitored how the RMSD and INF changed over the course of the simulation (Fig. 2c-f). We examined the furthest excursion from the reference conformation, as measured by the maximum RMSD (RMSD_max_) and minimum INF (INF_min_), and compared to the closest RNAnneal conformation, as measured by RMSD_min_ and INF_max_. The median RMSD_max_ among all reference structures was 6.1Å and the median INF_min_ was 0.85; RNAnneal sampled similar conformations, with a median RMSD_min_ of 7.4Å and a median INF_max_ of 0.80 across the set (Fig. 2c-d).

Ten of the ERCs contain PKs, which are excluded from the secondary structure prediction algorithms we employ. Consequently, the performance of RNAnneal’s candidate ensemble degrades as the number of PK base pairs in the ERC increases (superscript in Fig. 2e-f). Only one PK-containing ERC (6TFF^27^) was adequately sampled by RNAnneal (RMSD *<* 5Å and INF *>* 0.85) due to conformational sampling provided by FARFAR2 and MD. RNAnneal’s sampling was more consistent for the PK-free RNAs, with 5/6 ERCs (2MXS^28^, 6N2V^29^, 4RUM^30^, 6CB3^31^, and 6QN3^32^) being reproduced.

A Wilcoxon signed-rank test was employed to evaluate the statistical significance of the difference in means between RMSD_max_ and RMSD_min_ (and INF_min_ and INF_max_), which yielded an ambiguous *p*-value of 0.08 (0.13 for the INFs). A clearer picture arises when comparing the RMSD and INF values separately for the PK-containing and PK-free ERCs. Here, the test yielded a *p*-value of 0.01 for the RMSDs (0.02 for the INFs) among the PK ERCs, and a value of 0.30 (0.63 for the INFs) for the PK-free ones. Based on these results, we concluded that PK-free ERCs were generally modeled well by RNAnneal, while PK-containing ERCs were not, highlighting the ongoing challenges of modeling PKs.

Correlation between INF and RMSD values of RNAnneal conformations, computed separately for each riboswitch, yielded an *r* of − 0.49 ± 0.22 (*µ* ± *σ*) which was indicative of weak correlation (Fig. S14). This reflects the need for both alignment-based and alignment-free measures of structural fidelity for dynamic molecules such as RNA. Indeed, some ERCs exhibited large changes in RMSD during simulation while still maintaining the interaction network (4RUM^30^ and 6VMY) and vice versa (6E1W^33^). The INF was observed to be more robust to variations in structure caused by thermal fluctuations compared to the RMSD, which is consistent with previous work^26^. Notably, the non-WC interaction network was found to be highly dynamic for several of the ERCs (Fig. S5).

### The RNAnneal score is selective for experimentally resolved conformations

The performance of the RNAnneal score was bench-marked on the task of classifying ERCs and decoy conformations, with comparisons to two other state-of-the-art scores: the Rosetta score and DES-AMBER force field energy (see FARFAR2 and Molecular Dynamics). For this evaluation, we create a new set of structures by adding conformations from the reference simulations to the pool of conformations sampled by RNAnneal, and then ordering the set based on their scores (Fig. S13).

Classification performance was quantified by measuring how well each score prioritized near-ERCs (INF ≥ 0.85) over decoys (INF *<* 0.85) using two metrics: Area Under the Accumulation Curve (AUAC; see AUAC) and Boltzmann-Enhanced Discrimination of Receiver Operator Characteristic (BEDROC; see BEDROC).^34^ The AUAC is dominated by contributions from the lowest-ranked models, while BEDROC emphasizes classification accuracy among the highest-ranked structures via a tunable parameter *α* (equal to 8). Both metrics vary from zero to one, with the latter indicating perfect classification of ERCs over decoys. The RNAnneal score broadly outperformed the Rosetta and DES-AMBER scores on the classification task in both the AUAC and BEDROC metrics (Fig. 2g-h). RNAnneal had a median AUAC of 0.83 which was greater than the 0.67 for DES-AMBER or 0.59 for Rosetta. Similarly, the median BEDROC for RNAnneal was 0.40 compared to 0.26 for DES-AMBER or 0.16 for Rosetta. Statistical significance was assessed using the Wilcoxon signed-rank test, which yielded *p*-values of 0.02 (AUAC) and 0.02 (BEDROC) when comparing the Rosetta score to DES-AMBER, 0.008 and 0.03 for RNAnneal and DES-AMBER, and 0.004 and 0.008 for RNAnneal and Rosetta. These results provide evidence that the RNAnneal score differs from the Rosetta and DES-AMBER scores at a significance level of 0.05.

There was little variation in AUAC or BEDROC between different features and projections; instead, most of the variation is due to the latent dimension, *d* (Fig. S15). Similar results were obtained when RMSD ≤ 5Å is used as a classification criteria for defining near-ERCs (Fig. S16). Interestingly, when the TM models performed well on average, we observed an effect where the ensemble-averaged RNAnneal score had superior classification accuracy to the average, and sometimes even the best model (Fig. 2g-h).

Although PK-containing ERCs were not well-sampled by RNAnneal, the RNAnneal score was able to classify them apart from decoys. The ERCs with the five highest RNAnneal AUAC and BEDROC values (6DLQ^35^, 6Q57^36^, 6TFF^27^, 6UBU^37^, and 6WJR^38^) all contained PK base pairs (Fig. 2g-h). Still, not all PK-containing ERCs were successfully predicted by the RNAnneal score at the temperature used for the main results (*T* = 1); some ERCs were better prioritized at lower or higher temperatures (Fig. S17).

### Annealing improves classification and ranking of experimentally resolved conformations

RNAnneal encodes the scale of the energy landscape via a temperature parameter in the TM algorithm (see Thermodynamic Maps). The structure of the thermodynamic map reflects the following intutition: conformations that are low-lying on the energy landscape are sampled at low temperatures, and therefore ought to be mapped onto a low-variance prior, while those which are high on the energy landscape and sampled at hotter temperatures should be mapped onto a high-variance prior. To prioritize low-energy conformations, one may evaluate the RNAnneal score with a low-variance generative prior (see RNAnneal Score). The main results are presented for *T* = 1.0, which corresponds to one standard deviation above the minimum potential energy value. However, there were multiple instances where altering the temperature value improved RNAnneal’s performance on the ERC classification task.

The AUAC and BEDROC values improved for 11/16 riboswitches as the temperature was annealed from *T* = 10 to *T* = 1 (Fig. S17). The AUAC and BEDROC plateau as the inference temperature was decreased further for eight riboswitches (2MXS^28^, 6C27^39^, 6DLQ^35^, 6N2V^29^, 6Q57^36^, 6TFF^27^, 6UBU^37^, and 6WJR^38^), while classification performance continued to improve down to *T* = 0.1 for three (6E1W^33^, 6QN3^32^, and 6VMY^40^). The AUAC and BEDROC values showed little response to annealing for two riboswitches (4NYA^41^ and 4YAZ^42^), and while performance degraded slightly for three others (6CB3^31^, 4RUM^30^, and 3Q3Z^43^), the overall change in classification performance was relatively small.

### State-informed prediction of conformational ensembles

As stated previously, RNA structure is best understood in terms of conformational ensembles consisting of weighted states and representative conformations. ERCs provide critical and informative snapshots but do not elucidate the full ensemble. In addition to sampling 3D structures, ensemble prediction requires state assignments and the selection of representatives–a process that often relies on intimate knowledge of the system in question and human intuition. RNAnneal automates the process of generating a conformational ensemble by ranking states predicted by SPIB based on their average RNAnneal score, and then selecting the highest-scoring conformation from within each state as a representative structure (see State Ranking).

We predict ten-state ensembles for each of the 16 riboswitches and measure the state population and deviation from the ERC (RMSD and INF) for each representative. Note that the combined decoy and ERC dataset from the AUAC and BEDROC evaluation is employed here so that the model has an opportunity to predict ERCs in the ensemble, even if they were not sampled by RNAnneal. Ultimately, we found that RNAnneal predicted near-ERC structures (RMSD *<* 5Å and INF *>* 0.85) as representatives for 5/16 riboswitches, and 11/16 had representatives that satisfy a more relaxed similarity criteria (RMSD *<* 10Å or INF *>* 0.80) to the ERC (Fig. 3a). The top five representative conformers for each riboswitch are reported in Fig. S20-23.

### Quantification of ensemble heterogeneity via per-nucleotide interaction entropy

The simulations of the riboswitch ERCs revealed their interaction networks to be dynamic, especially in terms of non-WC interaction patterns (Fig. S5). Consistent with this observation, the riboswitch ensembles predicted by RNAnneal contained heterogeneous structures spanning wide ranges of RMSD and INF values (Fig. 3a). To assess heterogeneity in nucleobase interaction patterns across an ensemble, we develop the per-nucleotide interaction entropy.

The interaction entropy for a given nucleotide in a structural ensemble is the Shannon entropy of its Leontis-Westhof base-base contact classifications (see Interaction Entropy). When the interaction entropy is zero, the corresponding nucleotide is found in the same base-base interaction across all conformations in the ensemble, and as the interaction entropy increases, the nucleobase participates in more varied interactions. When measured in bits, an interaction entropy equal to *H* for nucleotide *i* implies that *i* participates in 2^*H*^ interactions across the ensemble; based on this, we bin the interaction entropy around integer values (colors in Fig. 3b).

The distribution of interaction entropies for the riboswitch ensembles predicted by RNAnneal indicate that a few of the riboswitches have mostly well-determined interaction patterns (light and dark blue in Fig. 3b), but most were predicted to have many nucleotides with variable interaction patterns (yellow and red in Fig. 3b). To aid in visualization of heterogeneity, the interaction entropy may also be used to color each nucleotide using the ranges indicated in Fig. 3b, as is done in Fig. S20-23.

### Evaluation of RNAnneal on experimentally resolved *holo* riboswitch conformations

RNAnneal was evaluated on fourteen *holo* riboswitches representing ligand-bound states that span transition-metal and natural-ligand complexes. Across this diverse set, RNAnneal frequently recovered conformations resembling experimentally resolved structures.

RNAnneal successfully predicted near-ERC representative structures for the transition-metal riboswitches (4RUM^30^, 6CB3^31^, and 6N2V^29^). The interaction network of the ERC for the cobalt-binding riboswitch (4RUM^30^) was reproduced in representative conformations predicted by RNAnneal (INF of 0.82 and RMSD of 8.1Å in Fig. 3a). Predicted structures for the cadmium- and manganese-binding riboswitches (6CB3^31^ and 6N2V^29^, respectively) were slightly less accurate in terms of INF (0.80 for 6CB3^31^ and 0.78 for 6N2V^29^) and RMSD (10.6Å for 6CB3^31^ and 8.5Å for 6N2V^29^). All three riboswitches had regions were the interaction entropy was low, indicating consistent structure across the ensemble (Fig. 3b). Visual inspection of the ERCs colored by the RNAnneal interaction entropy reveals that nucleobases with variable interaction patterns are in close proximity to bound transition metals in the *holo* structure (Fig. S20-23).

The rest of the *holo* ERCs bind natural ligands, which present a greater variety of interaction modes due to their diverse chemistry and spatially extended nature. RNAnneal recovered near-ERCs as representatives for four of the riboswitches, as quantified by an INF value greater than 0.85 and RMSD less than 5Å (3Q3Z^43^, 6TFF^27^, 6UBU^37^, and 6DLQ^35^). Superimposing the RNAnneal interaction entropy on each of the ERCs reveals variable interaction patterns for nucleobases close to the binding site(s) (Fig. 3c and S20-23). The next grouping includes two riboswitches (2MXS^28^ and 6QN3^32^) for which the RNAnneal representatives had moderate accuracy with respect to the ERC (0.7 *<* INF *<* 0.85 and 5Å *<* RMSD *<* 10Å). The INF for the neomycin sensing riboswitch (2MXS^28^) is low due to the relatively small number of interactions present; RNAn-neal predicted multiple conformations with RMSDs between 5 and 10Å, indicating good prediction accuracy. The glutamine II riboswitch (6QN3^32^) was solved via X-ray crystallography as a dimer, but was predicted as a monomer by RNAnneal. As a result, a large fraction of the bases in the ERC appear to be unpaired but actually form inter-chain interactions. Nonetheless, RNAn-neal predicts a representative conformation less than 10Å in RMSD from the ERC. Remarkably few nucleobases had a consistent interaction pattern, indicating that the monomer may not have a dominant stable fold (Fig. 3b and S20-23).

RNAnneal successfully recovered near-ERC representatives for 9/14 holo riboswitches. The remaining 5 systems (4NYA^41^, 4YAZ^42^, 6E1W^33^, 6Q57^36^, 6VMY^40^) showed lower agreement with the ERCs, as quantified by an INF value less than 0.7 and a RMSD greater than 10Å (Fig. 3a). Compared to the previous groupings, these riboswitches were predicted to have many nucleotides with high interaction entropy, consistent with a more flexible or dynamic structural ensemble (Fig. 3b). In general, nucleobases in close proximity to ligand binding sites have higher interaction entropies (Fig. S20-23).

### Evaluation of RNAnneal on experimentally resolved *apo* riboswitch conformations

Riboswitch ERCs are typically representative of the *holo* state due to increased conformational flexibility (i.e., heterogeneity) in the absence of a ligand. Consequently, the *apo* conformations of riboswitches are generally poorly characterized. Fortunately, two riboswitch ERCs have been solved in the *apo* state (6C27^39^ and 6WJR^38^). For the S-adenosyl methionine (SAM) III riboswitch (6C27^39^), multiple representative conformers with INF greater than 0.8 and RMSD between 10 and 15Å were predicted by RNAnneal (Fig. 3a). Similarly, RNAnneal predicted a representative conformation less than 5Å in RMSD from the ERC for the flavin mononucleotide (FMN) riboswitch in the apo state (6WJR^38^). The RNAnneal ensemble for SAM III riboswitch is somewhat homogenous as indicated by lower interaction entropies; the FMN riboswitch however, is more heterogeneous, likely due to PK base pairs present in the ERC which were poorly modeled during the structure sampling step (Fig. 3b)

### *De novo* prediction of *apo* riboswitch conformations

While experimentally resolved *apo* riboswitch structures are valuable benchmarks, most systems lack such reference structures, necessitating a *de novo* approach to characterize their conformational landscapes. RNAnneal is uniquely suited for this task since it is an unsupervised model.

We focus on the NAD^+^ riboswitch as a representative example and note that similar reasoning may be applied to other RNAs. This choice is motivated by three factors: the *holo* ERC (6TFF^27^; Fig. 3c) was predicted *de novo* by RNAnneal (Fig. 2e–f), found to have a fragile interaction network (Fig. S5), and was prioritized by the RNAnneal score (Fig. 2g–h).

Inspection of the location of the ten representative conformations in the SPIB latent space associated with the NAD^+^ riboswitch generated using RNAanneal reveals only five distinct clusters. The clusters are represented among the top five (of ten total) representative conformations (Fig. 3d). The first- and second-ranked states span the first information bottleneck (IB) component, suggesting that they are endpoints of the slowest structural transition. The representative conformation for the top-ranked state forms a single continuous helix with a six nucleotide overhang, while the second-ranked state forms a cloverleaf structure (Fig. 3d). The third-, fourth-, and tenth-ranked states cluster together and exhibit an internal loop that propagates up the 5’ side of the helix as the 5’ and 3’ ends pair together. The internal loop moves further up the helix in the fifth-through ninth-ranked states, where the representatives closely model the holo ERCs (both with and without the PK contacts). The transition from the *holo* ERC to the second-ranked state is unclear, but one may reasonably infer that the transition involves the loop propagating further up the helix.

## Discussion

RNAnneal introduces an unsupervised, *ab initio* framework for predicting three-dimensional structural ensembles of RNA. The strong performance of RNAn-neal on RNA structure prediction tasks may be attributed to multiple factors, including a hierarchical approach to sampling the structural ensemble, geometric and dynamics-based structure featurization, and deep learning models that incorporate physics-inspired inductive biases. The high performance of RNAanneal on structurally challenging systems indicates that that RNAnneal will be an essential tool for rapidly predicting three-dimensional conformations of RNA molecules.

Determining how to venture beyond experimental structures of both RNAs and proteins is critical in the post-AlphaFold2 world. RNAnneal generalizes beyond known ERCs by employing *ab initio* sampling to generate training data, and unsupervised deep learning to distinguish between experimentally resolved and decoy structures. Although in this manuscript RNAnneal has been benchmarked exclusively on the challenging case of riboswitches, the method is expected to generalize well since the structure generation pipeline relies exclusively on minimally parameterized physical models which have been previously evaluated on a variety of systems.^19,44^

RNAs are dynamic, flexible entities containing both persistently structured elements such as helices and dynamic elements such as internal loops. Variability of these structures is captured by RNAnneal’s ensemble output, and flexibility hotspots are conveyed by the interaction entropy. One may examine the riboswitch ERCs colored by their predicted interaction entropies in Fig. S20–23. Visual inspection of the structure datasets reveals that regions of moderate-to-high interaction entropy (yellow and red nucleotides) frequently coincide with the presence of stabilizing molecules such as metabolite ligands (e.g., Neomycin in 2MXS^28^ or PreQ1 in 6E1W^33^), metal ions (e.g., Mn^2+^ in 6N2V^29^ or Cd^2+^ in 6CB3^31^), or crystallization additives such as cobalt hexammine (e.g., in 6C27^39^ or 6UBU^37^). Regions with high interaction entropy may be important for ligand binding, allosteric regulation, or functional switching.

While RNAnneal is already demonstrated to be capable of tertiary structure ensemble prediction for different problems, it is a general and highly modular framework with many further directions for improvement. First, magnesium ions that facilitate folding by both neutralizing the phosphodiester backbone and binding to specific sites in folded RNA^45^, are not explicitly treated by RNAnneal. Future work should focus on both more appropriately modeling Mg^2+^ implicitly and predicting the location of explicit, bound Mg^2+^.^46^Second, employing templates for hard-to-sample motifs such as kink turns and multi-helix junctions^47–49^ may improve the quality of the sampled 3D structures, and would not bias the RNAnneal score since it is both unsupervised and parameterized *ab initio*. Third, the RNAnneal score may be extended to environmental variables like pressure and chemical potential, which have already been incorporated into the Thermodynamic Maps framework.^14^ Finally, the coverage of the candidate structure pool could be improved by including outputs from PK-aware secondary structure prediction algorithms^50–54^ and candidates models from physics-based^53–56^ and deep-learning^57–60^ tertiary structure prediction methods.

Overall, RNAnneal provides a target-specific method to generate and score 3D RNA structure ensembles from sequence alone that makes ensemble-level RNA structure prediction tractable and interpretable, and will help shine light on the sequence-structural ensemble relationships that underlie RNA diversity and function.

## Acknowledgments

This research was supported by the Intramural Research Programs of the National Institutes of Health, National Cancer Institute (NCI), Center for Cancer Research, Project BC011585 (to J.S.S.), by the National Institute of General Medical Sciences of the National Institutes of Health under Award Number R35GM142719, the National Science Foundation under Grant No. CHE-2044165 and the Maryland Technology Development Corporation (TEDCO) through the Maryland Innovation Initiative (MII) program under Grant No. 0725-0192. The authors thank UMD HPC’s Zaratan and NSF ACCESS (project CHE180027P) for computational resources. P.T. is an investigator at the University of Maryland-Institute for Health Computing, which is supported by funding from Montgomery County, Maryland and The University of Maryland Strategic Partnership: MPowering the State, a formal collaboration between the University of Maryland, College Park, and the University of Maryland, Baltimore. The content is solely the responsibility of the authors and does not represent the official views of the National Institutes of Health.

## Code Availability

An academic web server implementing RNAnneal is available for non-commercial academic use; access can be requested via https://go.umd.edu/rnanneal.

## Methods

### Secondary Structure Prediction

The secondary structure of an RNA is a simplified representation that specifies Watson-Crick base pairing. There are numerous, long-standing approaches available to predict secondary structures from the primary sequence.^61^ RNAnneal relies on three programs to predict RNA secondary structures: EternaFold^16^, LinearFold^17^, and LinearAliFold^17^. EternaFold is a probabilistic method for secondary structure prediction that has been benchmarked against several widely used secondary structure prediction algorithms (Algorithm 1). LinearFold is an algorithm for predicting secondary structures that has linear runtime with sequence length (Algorithm 2). LinearAliFold extends the functionality of LinearFold by implementing stochastic sampling (Algorithm 3). After a large number of secondary structures have been predicted by the above methods, the structures are aggregated and redundant ones are filtered out. Fig. S6 measures the fraction of correctly predicted bases for the predicted distributions of secondary structures for each riboswitch.

### FARFAR2

Rosetta’s Fragment Assembly of RNA with Full-Atom Refinement (FARFAR) is a hybrid knowledge- and physics-based approach to sampling models of 3D RNA structure.^62^ FARFAR samples 3D models by performing Monte Carlo sampling on a low-resolution potential using a library of experimentally validated structure fragments, and then relaxes the structure in a higher-resolution all-atom potential. The quality of the conformations is evaluated by the Rosetta score. FARFAR2 further improves the performance of FAR-FAR with updates to the sampling algorithms, scoring weights, and fragment libraries.^19^ The Rosetta score referred to in the main text and Fig. 2g-h corresponds to the `rna hires.wts` file in Rosetta v2021.16.61629. Algorithm 4 describes how RNAnneal uses FARFAR2 to sample a candidate model of the 3D structure with the primary sequence and a secondary structure as input.

### Clustering

A crucial step in RNAnneal is selecting representative conformations to simulate (see Molecular Dynamics) from the set of structures predicted by FARFAR2. To sample diverse conformations, we first compute G-vectors for the FARFAR2 conformations (see R- and G-vectors), and then project the G-vectors onto *d* = 2, 3, …, 26 principal components. For each *d* value, the data is clustered using the advanced density peaks algorithm^63^ implemented in `dadapy`^64^ with *Z* = 3. Representatives are selected based on the the *d* value that yields the maximum number of clusters (Fig. S7). To select, e.g., 200 candidate structures, we first sort the conformations within each cluster based on their Rosetta score and then sort the clusters based on the top-scoring conformation in each cluster. The clusters are then traversed in order, and the top-scoring conformation is selected as a representative and subsequently removed from the cluster. If there are fewer than 200 candidate structures after all clusters have been traversed, then they are reordered based on the new top-scoring conformation and the selection procedure is repeated until 200 conformations are obtained. The entire procedure is detailed in Algorithm 6.

### Molecular Dynamics

RNAnneal simulates models of 3D RNA structure with implicit solvent MD in order to introduce variations in structure caused by the thermal environment.

The simulations are conducted under the NVT ensemble using the OpenMM simulation engine and the DES-AMBER RNA force field.^44,65^ The conformations are equilibrated by gradually increasing the temperature from 1K to 300K and then increasing the integration timestep from 1fs to the production integration timestep of 5fs (Table II). The production simulation is 100ns and frames are saved at 1ns intervals; additional simulation parameters like ion concentration and solvent model are provided in Table III. The DES-AMBER score is the potential energy of the MD simulations (i.e., the energy values of the force field and solvent model) standardized by first aggregating the energy across all simulations (for a given RNA) and then subtracting the minimum value and dividing the standard deviation.

**TABLE 2.** Molecular Dynamics Equilibration Procedure.

| Stage | Duration (ps) | Timestep (fs) | Temperature (K) |
| --- | --- | --- | --- |
| Equilibration 1 | 1 | 0.1 | 1 |
| Equilibration 2.1 | 1 | 1 | 10 |
| Equilibration 2.2 | 1 | 1 | 20 |
| Equilibration 2.3 | 1 | 1 | 30 |
| Equilibration 2.4 | 1 | 1 | 40 |
| Equilibration 2.5 | 1 | 1 | 50 |
| Equilibration 3.1 | 2 | 1 | 100 |
| Equilibration 3.2 | 2 | 1 | 150 |
| Equilibration 3.3 | 2 | 1 | 200 |
| Equilibration 3.4 | 2 | 1 | 250 |
| Equilibration 3.5 | 2 | 1 | 300 |
| Equilibration 4.1 | 20 | 1 | 300 |
| Equilibration 4.2 | 20 | 2 | 300 |
| Equilibration 4.3 | 20 | 3 | 300 |
| Equilibration 4.4 | 20 | 4 | 300 |
| Equilibration 4.5 | 20 | 5 | 300 |
| Production | 100,000 | 5 | 300 |

**TABLE 3.**
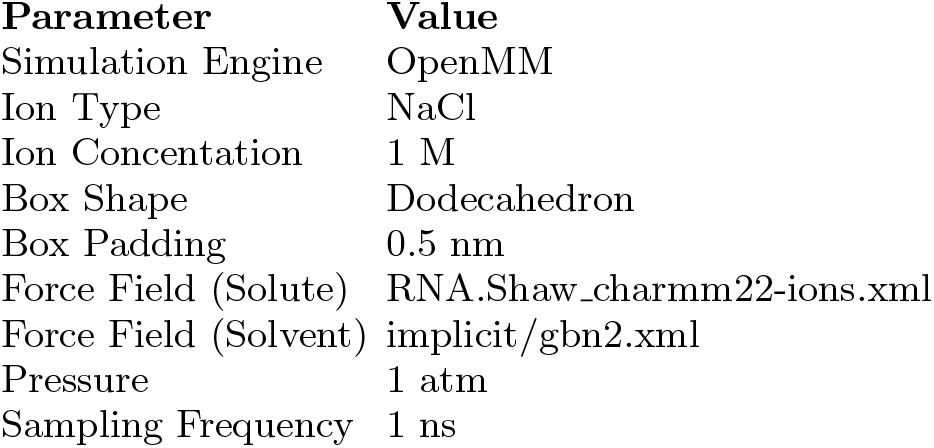
Molecular Dynamics Simulation Parameters.

| Parameter | Value |
| --- | --- |
| Simulation Engine | OpenMM |
| Ion Type | NaCl |
| Ion Concentration | 1 M |
| Box Shape | Dodecahedron |
| Box Padding | 0.5 nm |
| Force Field (Solute) | RNA.Shaw_charmm22-ions.xml |
| Force Field (Solvent) | implicit/gbn2.xml |
| Pressure | 1 atm |
| Sampling Frequency | 1 ns |

### R- and G-vectors

RNAnneal featurizes conformations with two nucleobase-centric representations constructed from R- and G-vectors.^66^ An R-vector is a displacement vector **r**_*ij*_ directed between the six-membered aromatic ring of nucleobases *i* and *j*. The R-representation of a conformation with *N* nucleobases is thus a tensor of shape *N* × *N* × 3. Similarly, the G-representation is a *N* × *N* × 4 tensor consisting of G-vectors that capture the geometry of localized base-pairing and stacking interactions. The G-vector, **G**_*ij*_, is computed as

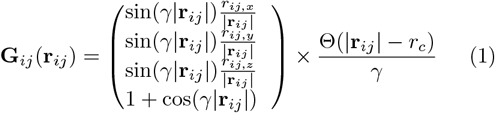

where Θ is a step function, *r*_*c*_ is a cutoff of 2.4Å, and *γ* = *π/r*_*c*_.

### SPIB

The State Predictive Information Bottleneck (SPIB) framework allows one to embed a high-dimensional dynamical system into a lower-dimensional latent space while preserving the slowest degrees of freedom of the system.^20^ SPIB takes the form of a time-lagged autoen-coder, where high dimensional coordinates at time *t* are passed to the encoder and embedded in a latent space; the decoder must then predict the state label at time *t* + Δ*t* solely from the latent embedding. SPIB has two main hyperparameters: the time lag, Δ*t* and a regularizer, *β*. The time lag corresponds to the lifetime of the predicted states, while *β* regularizes the latent space to follow a mixture of Gaussians prior. We train two sets of SPIB models on the first 8 principal components of the R- and G-vectors. For each feature type, we train seven models with the number of IB components, *d*, ranging from 2 to 8. For all models, we use a linear encoder, non-linear decoder, a 2ns lag time, and set *β* = 0.05. The SPIB implementation is provided by the `af2rave` python package.^67^

### Thermodynamic Maps

A thermodynamic map (TM) is a diffusion model variant that models the dependence of a structural ensemble on physical parameters like temperature, pressure, or chemical potential by representing their statistical effect on the prior distribution.^12,14,15^ Temperature, for example, is represented by tempering the prior (i.e., adjusting the variance) on a per-sample basis, while pressure and chemical potential correspond to exponential tilting operations that shift the mean of the prior.^12,14^ Once trained, the TM model can then be used to generate new samples or evaluate the likelihood of existing samples at a given set of thermodynamic conditions. In this work we modify the construction so that the prior is tempered based on the energy of the conformations. Concretely, for a given representation **x**_0_, we set the variance of the prior equal to the DES-AMBER score, *U*_0_, of the conformation corresponding to **x**_0_ (see Molecular Dynamics; Fig. 1c). The RNAnneal Score for a given conformation is computed from the log-likelihoods of the latent representations under the trained models. The log-likelihood calculation is summarized in Algorithm 7 and described in detail in Ref.^68^ Our TM implementation infers the energy-dependence of the conformational ensemble, and as a result one must choose a variance (i.e., a temperature) for the prior when computing the score. All the results in the main text employ a unit variance prior, but other values sometimes lead to better performance (Fig. S17-S19).

### RNAnneal Score

RNAnneal scores conformations using an ensemble of Thermodynamic Maps trained on geometric and energetic features extracted from a set of 3D conformations (Fig. 1b-d). A total of 28 TMs are trained for each RNA, covering all four combinations of representations (R- and G-vectors) and projections (PCA and SPIB). For each feature-projection combination, the number of projection components *d* varies from 2 to 8 inclusive. After training, the RNAnneal score is computed from the log-likelihoods of the training data under the TM ensemble (Algorithm 7 and 8).

### Reference Structures

RNAnneal’s structure prediction performance was evaluated on a set of riboswitch sequences with structures that have been solved by high-resolution experimental methods (Table I and Fig. S4). The reference ERCs were prepared by downloading the associated Protein Data Bank (PDB) entry, removing any non-nucleic residues, filling in missing nucleobases using FARFAR2, and subsequently minimizing the conformation with the Rosetta energy function (see Algorithm 5).

### RMSD

Root Mean Squared Deviation (RMSD) is a metric commonly used to evaluate discrepancies between structure models. We evaluate the heavy-atom RMSD of each model after alignment to the native structure. The RMSD between a model **x** and reference **y** conformation containing *N* heavy atoms is defined as

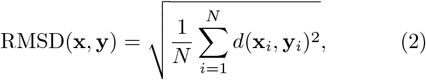

where *d*(**x**_*i*_, **y**_*i*_) measures the distance between the *i*-th atom in both models. The RMSD values for RNAnneal conformations and molecular dynamics simulations of the reference structure are reported in Fig. S9 and Fig. S8.

### INF

Interaction Network Fidelity (INF) quantifies the similarity between two conformations based on their Leontis-Westhof base pairing and stacking classifications.^26,69^ Let *S*_*m*_ and *S*_*r*_ denote the sets of nucleobase interactions (base pairs and base stacks) in the model and reference conformations, respectively. The INF is defined as the Fowlkes-Mallows Index between these interaction sets:

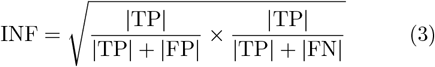

where TP = *S*_*m*_ ∩ *S*_*r*_ are interactions present in both structures, FP = *S*_*m*_ *S*_*r*_ are interactions present only in the model, and FN = *S*_*r*_ *S*_*m*_ are interactions present only in the reference. Minor, localized structural differences (e.g., at a helix junction) may yield large RMSD values due to poor global alignment. Because the INF compares interaction networks directly without requiring alignment, it is more robust to such variations and to thermal fluctuations. We compute the INF using the ClaRNA^70^ implementation in rna-tools^71^. INF values are reported in Fig. S5 and S11-12.

### AUAC

The Area Under the Accumulation Curve (AUAC) is a metric suited for evaluating classification task performance (Fig. 1e and 2g).^34^ Let *x* ∈ [0, 1] be the normalized rank and *f* : *r* → { 0, 1 } be an associated set of labels, with 1 indicating the positive (near-ERC) class and 0 indicating the decoy class. The empirical CDF is then

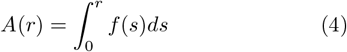

and the area under the accumulation curve is

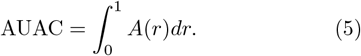

When the AUAC is 1, the classification is perfect, i.e., members of the positive class are always ranked above decoys. The accumulation curves for the 15 riboswitches investigated in the main text are provided in Fig. S18 and S19.

### BEDROC

The Boltzmann-Enhanced Discrimination of Receiver Operator Characteristic (BEDROC) modifies the Area Under the Accumulation Curve (AUAC) to focus on early recognition of the positive (near-ERC) class (Fig. 1e and 2g).^34^ The AUAC is skewed towards late recognition since contributions from the lowest ranked samples dominate the area calculation in Eq. 5. One may introduce an exponentially decreasing weight on the rank in the AUAC calculation to focus on early recognition, i.e.,

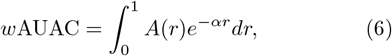

which is normalized to obtain

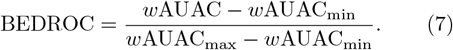

The tunable parameter *α* controls the focus on early recognition. Our choice of *α* = 8 is such that the top 20% of the normalized rank contributes 80% of the *w*AUAC. Early recognition is a suitable evaluation metric when there is a large class imbalance and accurate classification among the highest ranks is essential.

### Interaction Entropy

Interaction entropy measures how heterogeneous each nucleobase’s interaction patterns are across an ensemble of conformations. First, each nucleobase *i* is assigned a Leontis–Westhof (LW) base-base contact classification for every structure in the ensemble.^69^ The distribution of these classifications defines probabilities *p*_*i*_(*c*_*j*_) for each contact *c* with partner nucleotide *j*. The interaction entropy is then the Shannon entropy of this distribution,

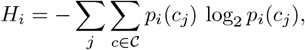

where 𝒞 denotes the set of LW classes. Expressing the Shannon entropy in bits lends a direct interpretation: a nucleotide with interaction entropy *H* participates in 2^*H*^ distinct base-base interactions across the ensemble (Fig. 3b). The probabilities may be computed as un-weighted frequencies or with prior weights associated to each structure. In this work we use state probabilities (see State Ranking) computed from the RNAnneal score as prior weights.

### State Ranking

We briefly describe the procedure by which a conformational ensemble, i.e., a small number of representative conformations and their weights, is predicted by RNAnneal. We project the RNAnneal conformations onto the *d* = 2 information-bottleneck (IB) space learned by SPIB to obtain latent representations { **x**_0_, **x**_1_, …, **x**_*N*_ }. We further assign the predicted state label, *y*_*i*_ ∈ { *y*_0_, *y*_1_, …, *y*_*M*_ } and RNAnneal Score, *s*_*i*_, to each conformation. Next, we histogram the conformations on a *K* × *L* grid in the IB space (Fig. 2j), using the mean *s* in each bin (denoted by 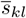) to define a Boltzmann-like weight 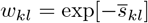. We assign each populated bin a state label, *y*_*kl*_, based on the most frequent state label of the conformations in the bin, and define a total weight, *W*_*i*_, for state *y*_*i*_ by 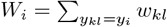. Finally, normalizing the state weights yields state probabilities *p*_*yi*_, and the frame with the minimal *s*_*i*_ within each state is taken as its representative conformation. The entire procedure is described in Algorithm 10. The states predicted in the main text (Fig. 3) and supplement (Fig. S20-S23) were predicted based on a *d* = 2 SPIB model trained on the first 2 PCA components of the G-vectors.

## Supplementary Information

### Algorithm 1

Sampling RNA secondary structures with `EternaFold`

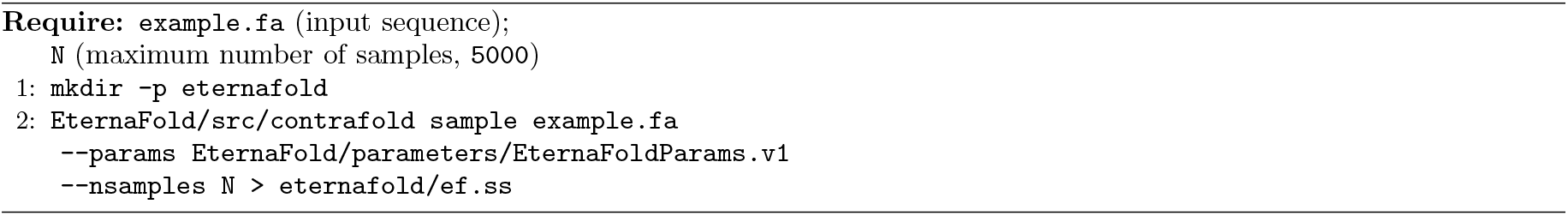

### Algorithm 2

Sampling RNA secondary structures with `LinearFold`

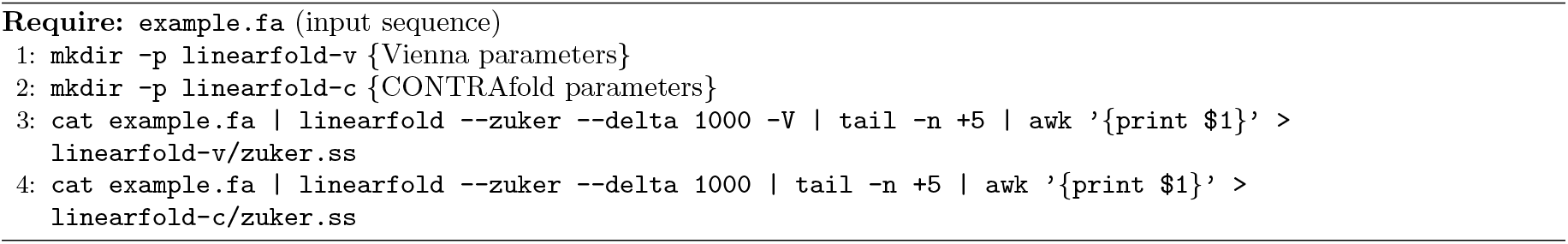

### Algorithm 3

Sampling RNA secondary structures with `LinearAliFold`

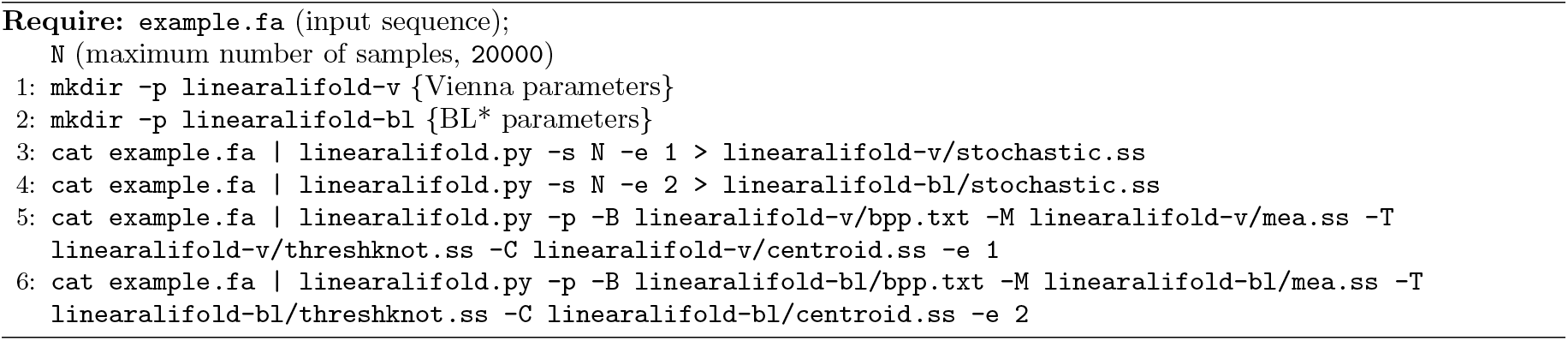

### Algorithm 4

Conformational sampling with FARFAR2

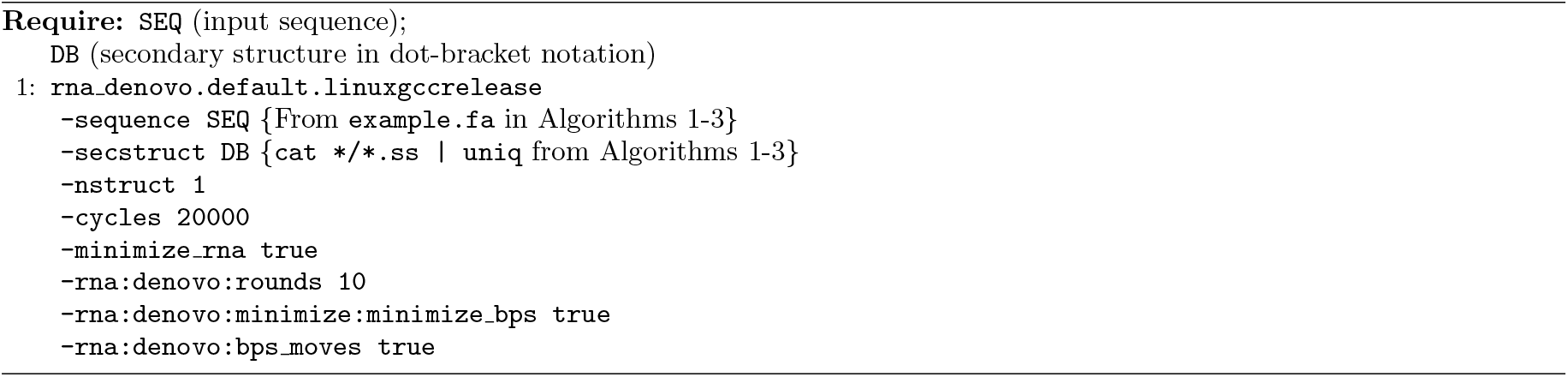

### Algorithm 5

Reference conformation preparation with FARFAR2

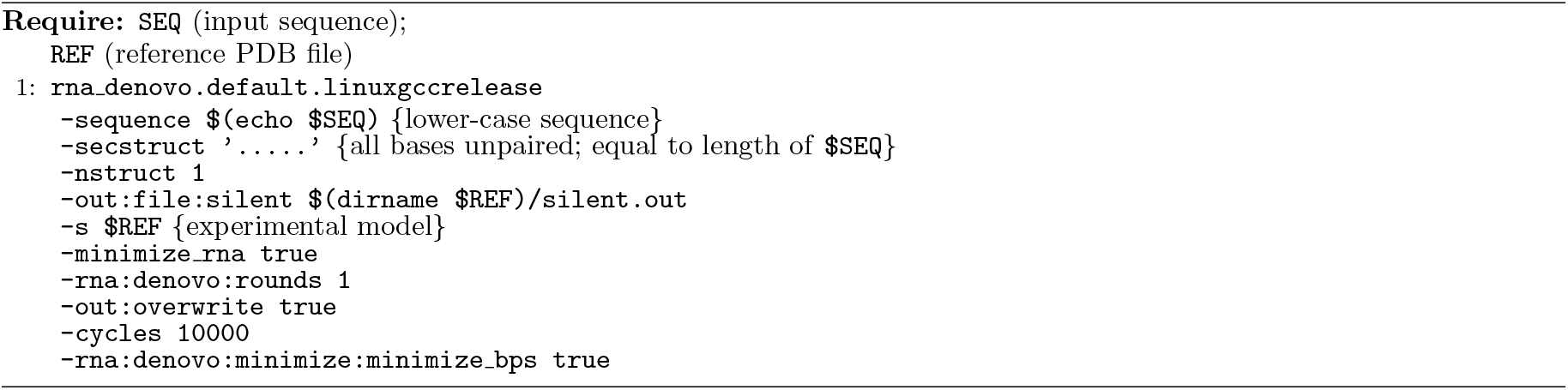

### Algorithm 6

Unsupervised selection of diverse candidate conformations

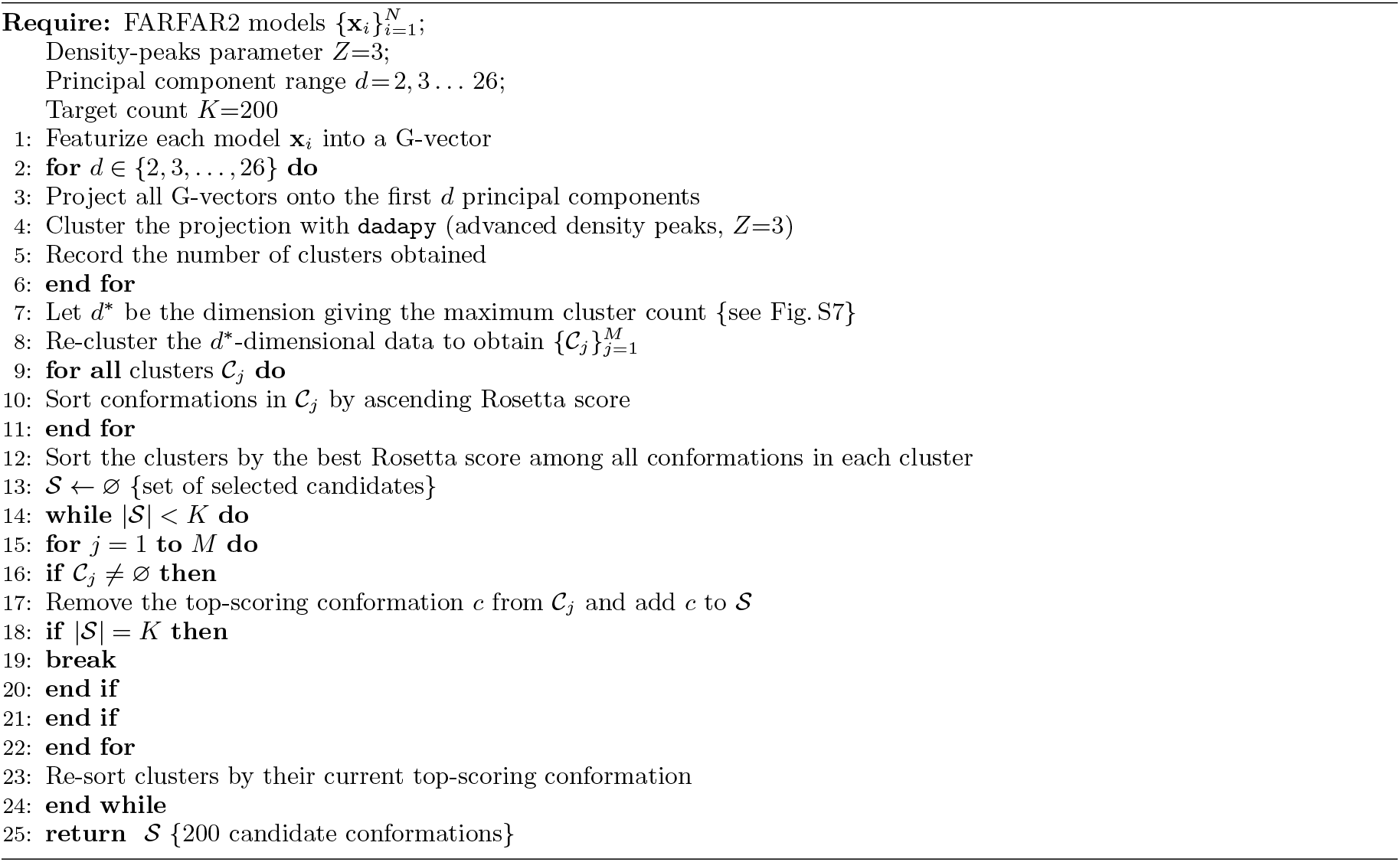

### Algorithm 7

Likelihood estimation for a thermodynamic map (diffusion model) using the Hutchinson trace estimator

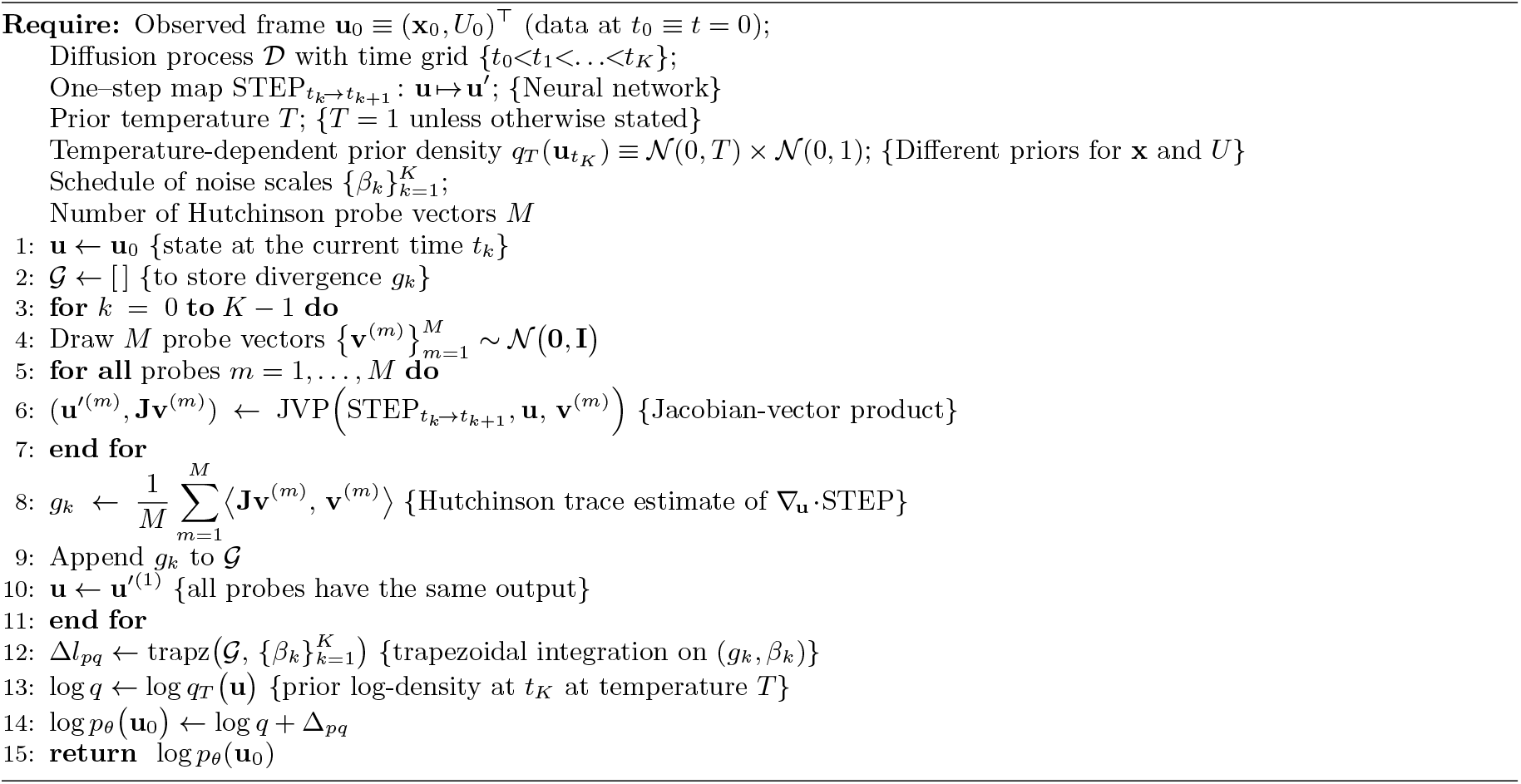

### Algorithm 8

RNAnneal Score

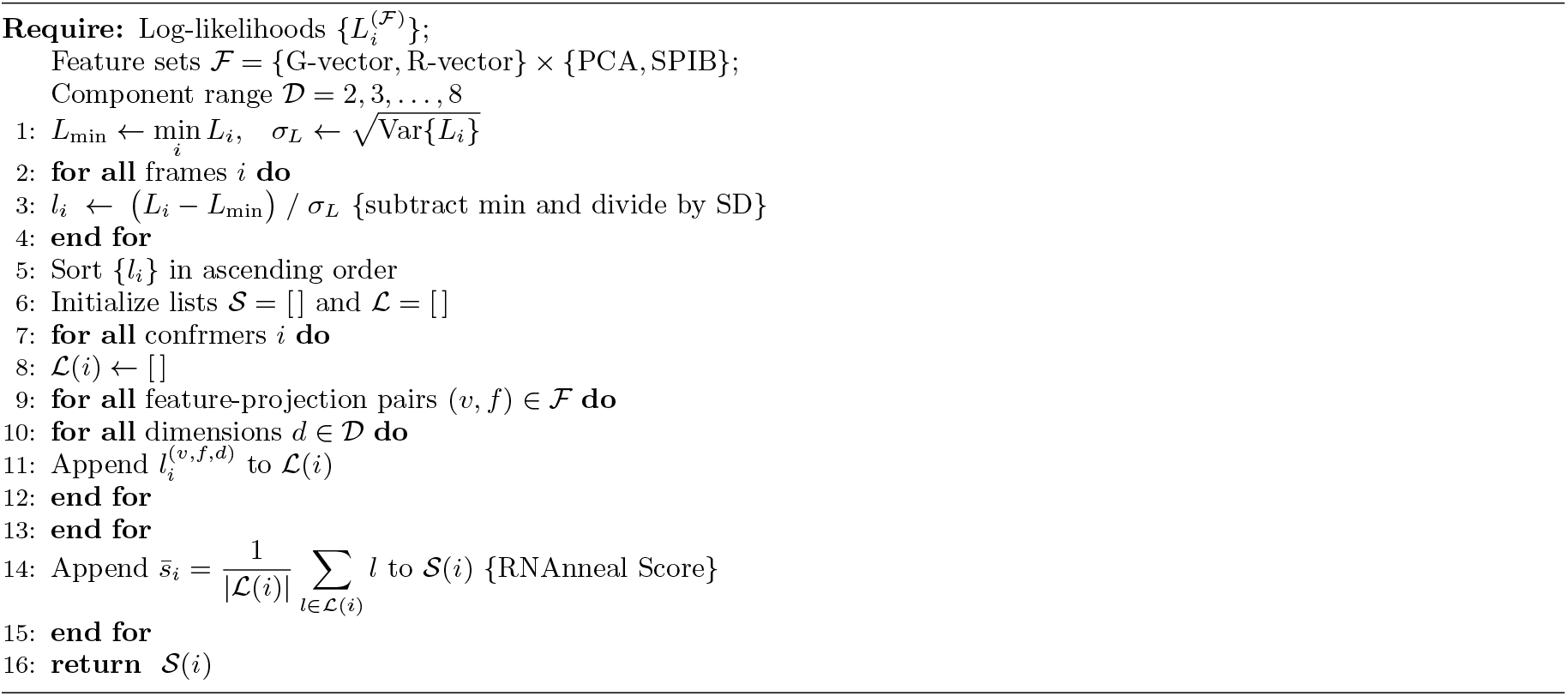

### Algorithm 9

Interaction Entropy calculation

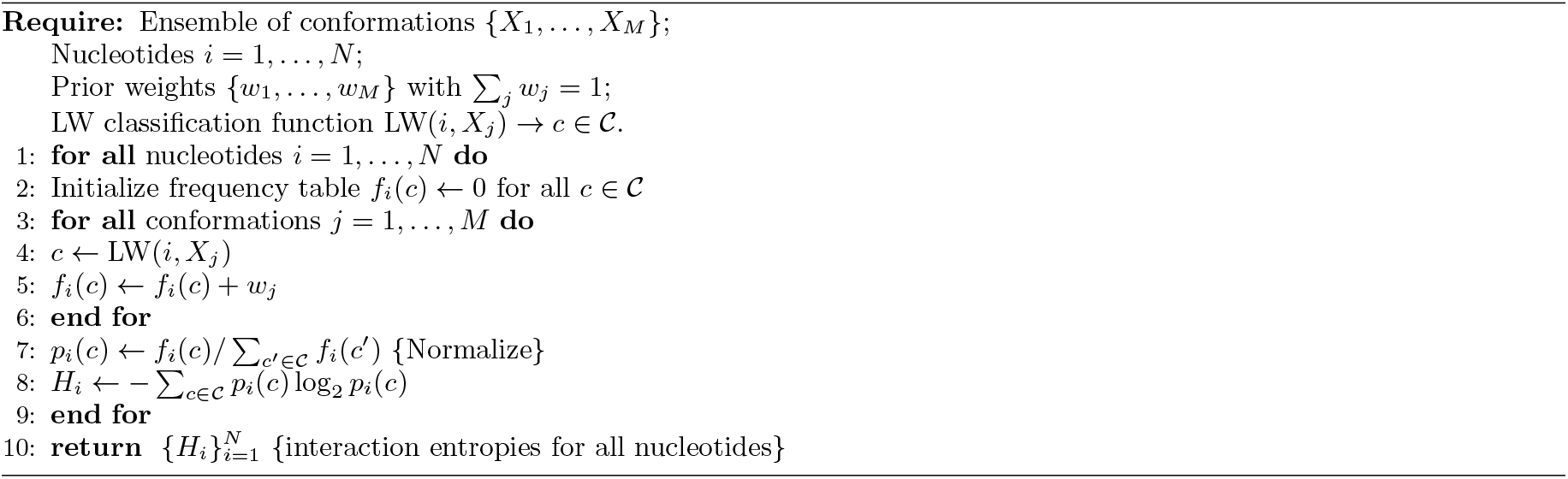

### Algorithm 10

Boltzmann-weighted state ranking and representative selection

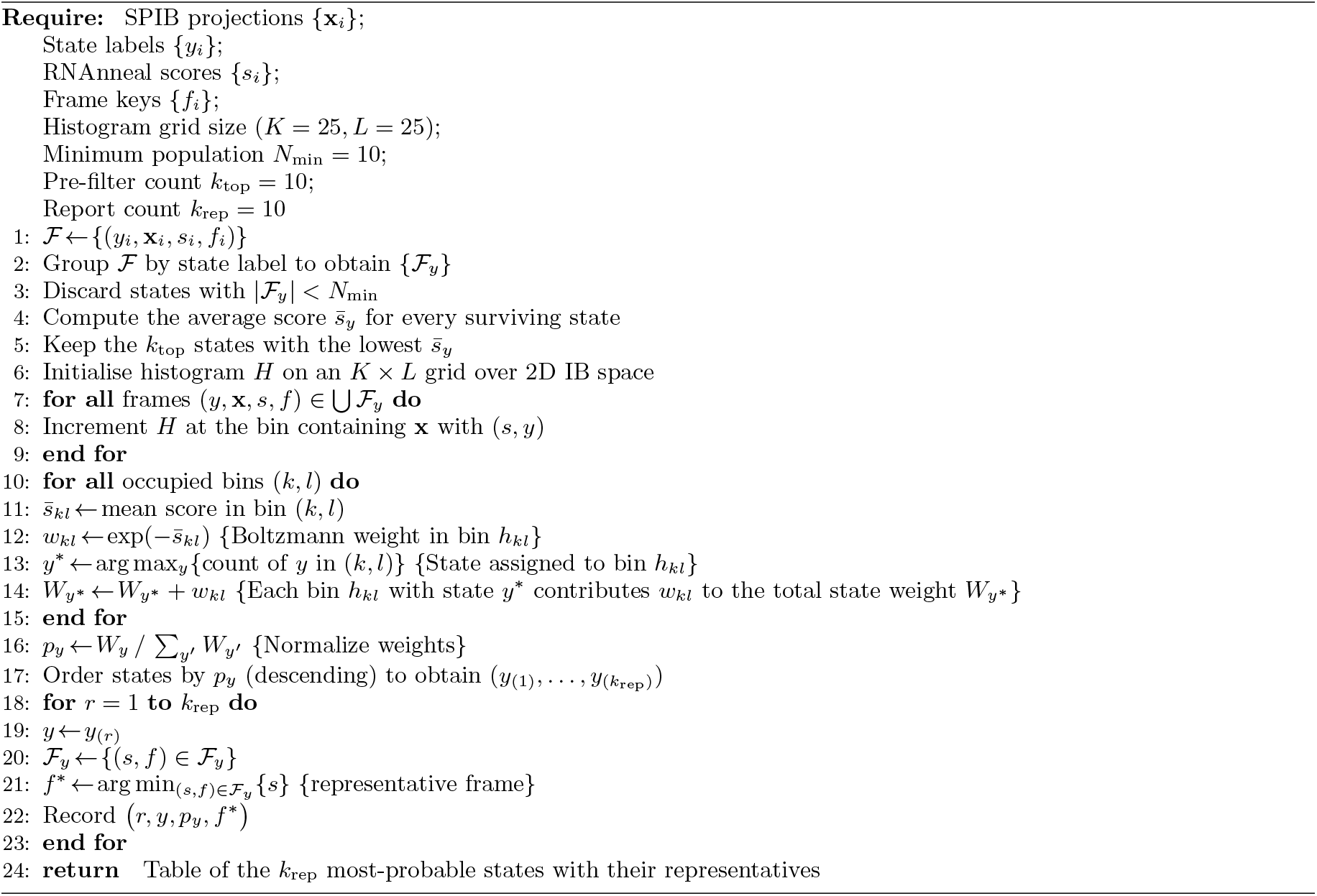

